# Biomimetic and Non-biomimetic Extraction of Motor Control Signals Through Matched Filtering of Neural Population Dynamics

**DOI:** 10.1101/023689

**Authors:** Islam S. Badreldin, Karim G. Oweiss

## Abstract

Brain-machine interfaces rely on extracting motor control signals from brain activity in real time to actuate external devices such as robotic limbs. Whereas biomimetic approaches to neural decoding use motor imagery/observation signals, non-biomimetic approaches assign an arbirary transformation that maps neural activity to motor control signals. In this work, we present a unified framework for the design of both biomimetic and non-biomimetic decoders based on kernel-based system identification. This framework seamlessly incorporates the neural population dynamics in the decoder design, is particularly robust even with short training data records, and results in decoders with small filter delays. The theory and results presented here provide a new formulation of optimal linear decoding, a formal method for designing non-biomimetic decoders, and a set of proposed metrics for assessing decoding performance from an online control perspective. The theoretical framework is also applicable to the design of closed-loop neural control schemes.

## 1 INTRODUCTION

The last decade witnessed tremendous advances in efferent brain-machine interface (BMI) control. At the core of BMI operation lies the so-called neural ‘decoder’ – a mathematical mapping that maps patterns of neural activity (input signals) into a ‘decoded’ signal (output signal) that is subsequently used to control external devices. Traditionally, decoders are designed such that the decode mimics actual kinematics (**Serruya et al.**, 2002; **Carmena et al.**, 2003; **Paninski et al.**, 2004; **Hochberg et al.**, 2006) or kinetics (**Fagg et al.**, 2009; **Song and Giszter**, 2011; **Suminski et al.**, 2013). In both cases, biomimetic decoder design requires a synchronized set of recorded neural activity and kinematic/kinetic correlates to estimate the filter functions. An alternative approach to neural decoder design is *non-biomimetic* decoding. Rooted in the pioneering work of **Fetz** (1969), non-biomimetic decoding directly assigns arbitrary mathematical mappings from the neural input signals to the decode via arbitrary filter functions. To date, non-biomimetic decoders rely on heuristics to design these filter functions. For example, exponential averaging (**Fetz**, 1969; **Fetz and Finocchio**, 1971; **Fetz and Baker**, 1973) and moving average filters (**Moritz et al.**, 2008; **Moritz and Fetz**, 2011; **Koralek et al.**, 2012; **Clancy et al.**, 2014) have been used for non-biomimetic decoding. Yet, the link between biomimetic and non-biomimetic decoder designs has been lacking.

Recently, we proposed a formal non-biomimetic decoder design method that rely on mathematical optimization (**Badreldin et al.**, 2013; **Badreldin and Oweiss**, 2014). In this work, we formalize the decoder design problem using the general class of kernel-based system identification methods. We first demonstrate the connection between neural decoding and system identification, and then we review results from the machine learning and systems identification literature (**Pillonetto et al.**, 2014) to link both biomimetic and non-biomimetic decoders from a ‘matched filter’ perspective that only differ in the objective function to be optimized.

## 2 THEORY AND METHODS

### 2.1 OPTIMAL LINEAR DECODER AND WIENER FILTERING

Optimal linear decoding has been widely used in the BMI community (**Warland et al.**, 1997; **Serruya et al.**, 2002; **Carmena et al.**, 2003; **Paninski et al.**, 2004; **Patil et al.**, 2004; **Hochberg et al.**, 2006; **Fagg et al.**, 2009; **Suminski et al.**, 2010). Neural spike trains from single-or mutli-unit activity are first converted to neural spike counts in predefined time bins to form a neural response time series for each individual unit, which is subsequently filtered (convolved) with linear filters of finite duration. In the final step, the unit filter outputs (referred to as unit outputs for short) are added together to obtain the ‘decode’, i.e. the quantity that is being decoded. These steps are pictorially illustrated in Figure 1.

**Figure 1.**
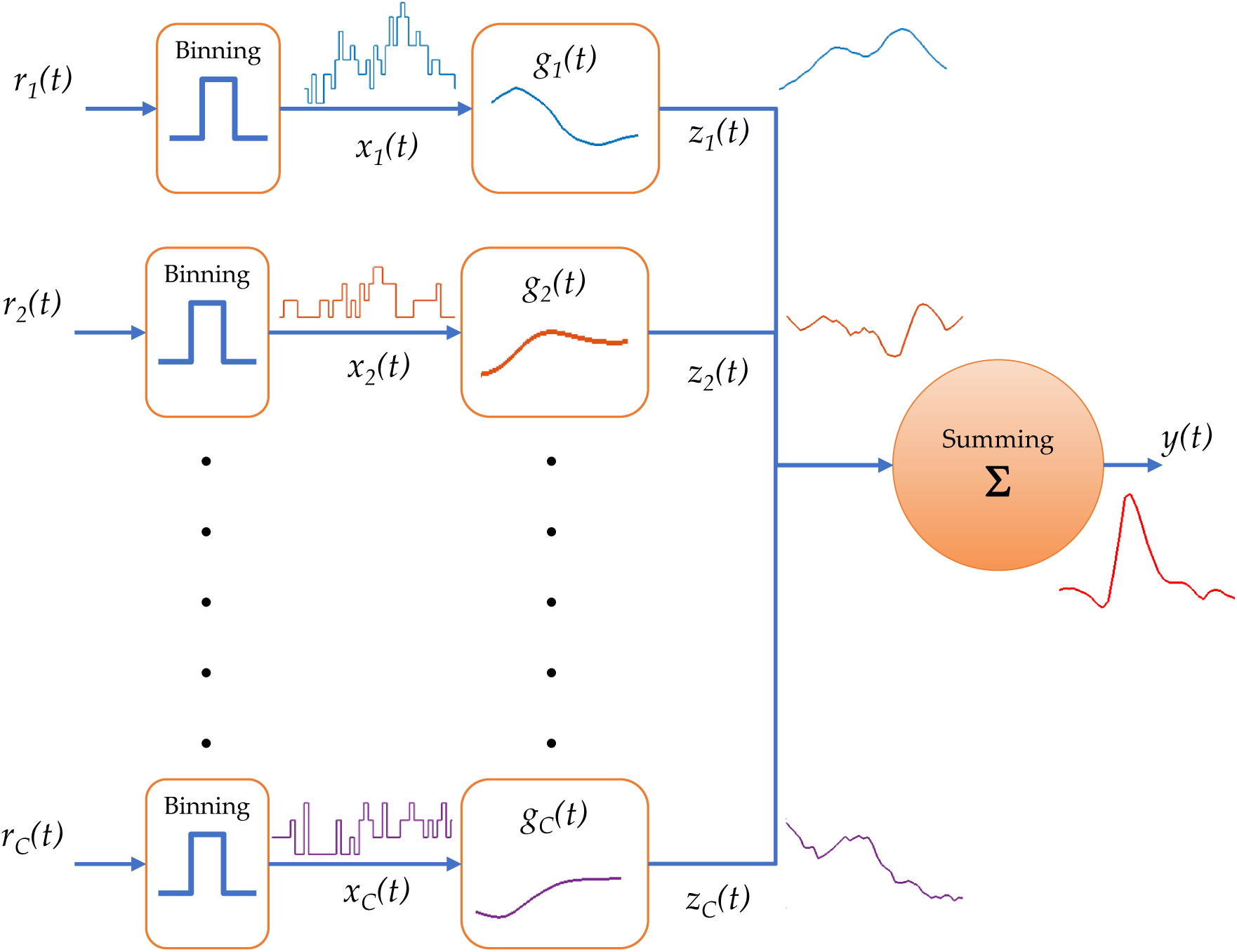
Linear decoder block diagram. Raw spike trains, *r*_*i*_(*t*), are counted using predefined time bins to produce binned spike counts, *x*_*i*_(*t*), which are subsequently filtered using a unit filter function, *g*_*i*_(*t*), to produce unit outputs, *z*_*i*_(*t*). The decode is computed as the summation of all unit outputs.

If we denote raw spike trains by *r*_*m*_(*t*), binned spike trains by *x*_*m*_(*t*), the unit filter coefficients as a function of time by *g*_*m*_(*t*), and the ‘decode’ by *y*(*t*), then the optimal linear decoder equation (**Warland et al.**, 1997; **Paninski et al.**, 2004) is given as^1^

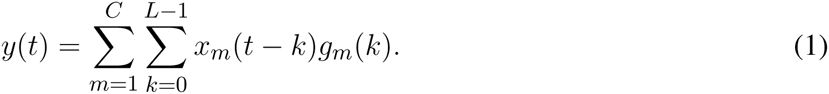

The inner sum is the filtering operation for binned spike trains from a particular unit to produce a unit output, and then the outer sum simply sums all unit outputs to produce the final ‘decode’. Given a synchronized set of input-output pairs {(*x*_*m*_(*t*), *y*(*t*)): *t* = 0, 1, 2, *…, T* 1; *m* = 1, 2, *…, C*}, the unit filter coefficients are solved for by rewriting Equation 1 in matrix form (**Paninski et al.**, 2004)

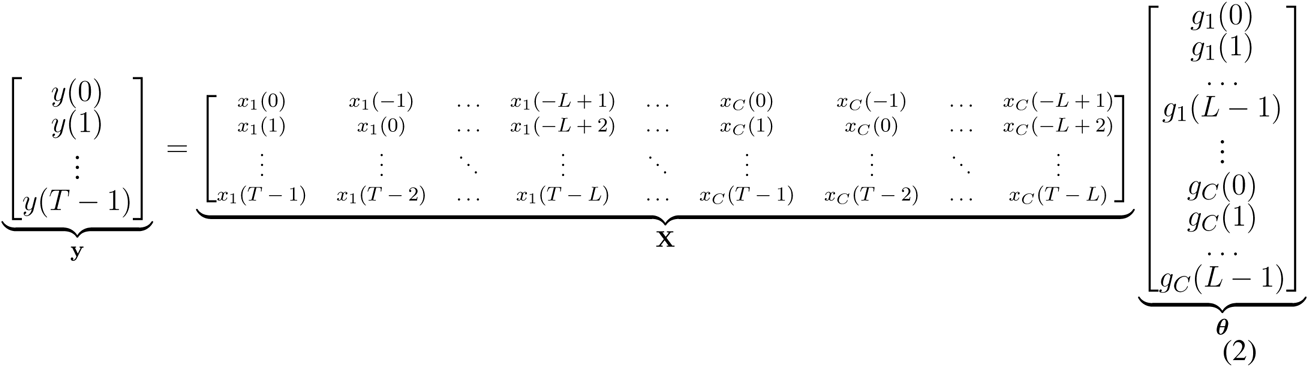

The solution is obtained by solving the so-called *normal equations*, **X**^T^**X *θ*** = **X**^T^**y**, and the ‘optimal linear filters’ are given succinctly by

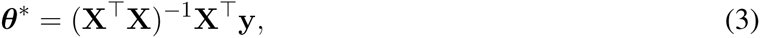

where the matrix 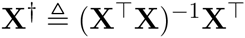 is known as the pseudo-inverse matrix of **X** (**Strang**, 2009).

Hence, by stacking all filter coefficients for all units {*g*_*m*_(*t*): *m* = 1, 2,…, *C*; *t* = 0, 1,…, *L* – 1} into a parameter vector ***θ*** and arranging the binned neural spike counts into an input data matrix **X**, we obtain the pseudo-inverse solution as in (3). According to the *projection theorem*, the pseudo-inverse solution can be interpreted as the solution that minimizes the error between the decode **ŷ**= **X *θ***, and the actual output **y** (**Strang**, 2009)

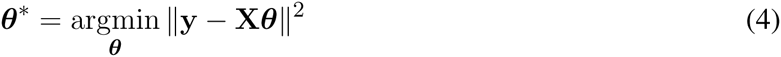

Hence, the decoder’s underlying generative data model is

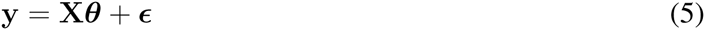

where ***ϵ*** = [*ϵ* (*t*): *t* = 0,…, *T* 1]^T^ is assumed to be white Gaussian noise. This assumption on the additive noise term makes the solution in (3) the *maximum likelihood* solution for the unknown decoder coefficients {*g*_*i*_(*t*): *i* = 1, 2,…, *C*; *t* = 0, 1,…, *L* - 1} (**Bishop**, 2006).

This optimal linear decoder solution is commonly referred to as finite duration Wiener filtering (**Carmena et al.**, 2003; **Sanchez et al.**, 2004; **Patil et al.**, 2004; **Kim et al.**, 2007; **Fagg et al.**, 2009; **Suminski et al.**, 2010), in which it is assumed that the input-output second order statistics are known (**Sanchez et al.**, 2005; **Kim et al.**, 2006). Given the *LC* × *LC* correlation matrix between the inputs as

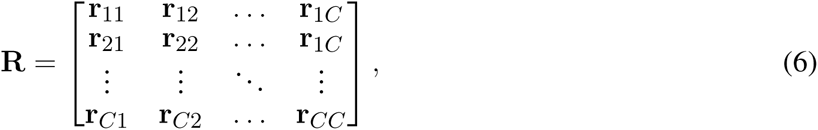

where **r**_*mn*_ is a *L* × *L* correlation matrix between units *m* and *n* at different time lags *L* (autocorrelation in case *m* = *n* and crosscorrelation in case *m* ≠ *n*), and the *LC* × 1 cross-correlation vector between the inputs and the output as

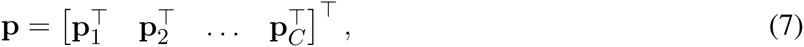

where **p**_*m*_ is a *L* × 1 cross-correlation vector between unit *m* and the output *y*(*t*) at different time lags *L*, the weights of the finite-duration Wiener filter are estimated using the Wiener-Hopf solution in the time domain (**Wiener**, 1949; **Kim et al.**, 2006)

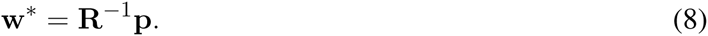

The intimate relationship between optimal linear decoding and Wiener filtering is seen when we consider the definitions of the correlations in **R** and **p** given by

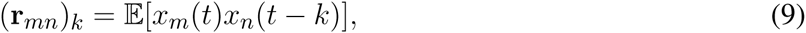

and

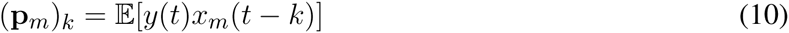

respectively. Thus, the solution given in (3) is the large-sample approximation of the idealized solution given in (8). In fact, the matrix

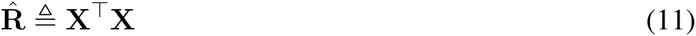

has its elements

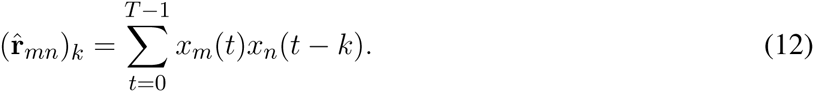

Therefore, the data-based matrix 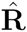 is an unnormalized estimate of the sample covariance matrix, which is a biased estimator of **R**. Similarly, the vector 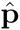 is an unnormalized biased estimate of **p**. Given a large data set for estimating the decoder, we expect the data-driven solution (3) to *converge* to the idealized solution (8). The problem that is faced in practice is: how *fast* is this convergence? It is widely reported that estimating Wiener-like filters requires about 10 minutes of data (**Carmena et al.**, 2003; **Paninski et al.**, 2004; **Patil et al.**, 2004; **Flint et al.**, 2012, 2013). The implications of using short finite-data samples to estimate the decoder filters are investigated below.

### 2.2 CONNECTION WITH SYSTEM IDENTIFICATION

The optimal linear filters structure presented in Figure 1 is a very general structure for modeling black-box input-output data. In the general case, the filter coefficients for each unit can be of infinite length. For example, the Kalman filter employed in the context of neural decoding gives exponentially decaying filter weights to all of the temporal history of the neural input, due to its recursive nature, as was formally derived in Appendix A.3 in **Wu et al.** (2006) (see also the linear dynamical system formulation by (**Gowda et al.**, 2012, 2014)). With unit filters of infinite duration, the linear decoding equation (compare to (1)) becomes

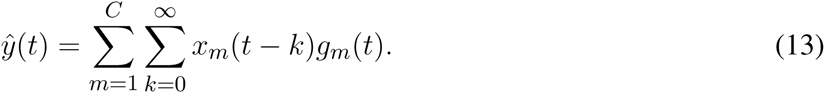

Hence, decoders with finite-duration linear filters^2^ are indeed *lower-order approximations* of general infinite-duration linear filters^3^ (**Ljung**, 1999). The goodness of these approximations depends on the dynamics of the approximated filters. Slower dynamics necessitate unit filters of longer duration to fully capture their temporal extent, whereas faster dynamics can be captured using unit filters of a shorter duration. In other words, the approximation error is bounded by the rate at which the individual unit filter coefficients decay to zero as a function of time (**Ljung**, 1999; **Wahlberg et al.**, 2005). The dynamics of the unit filter coefficients depend on the decode dynamics, where for the latter it is desirable to have a dynamical bandwidth on the order of 1-3 Hz in case the decode represents a command signal that directly drive a robot, i.e. a motor control signal (**Willett et al.**, 2013). Therefore, as widely reported, the unit filter length is expected to be around one second (**Serruya et al.**, 2002; **Carmena et al.**, 2003; **Hochberg et al.**, 2006; **Fagg et al.**, 2009; **Suminski et al.**, 2010).

### 2.3 SINGULAR VALUE DECOMPOSITION OF OPTIMAL LINEAR DECODERS

Consider the SVD of the neural data matrix **X**, which is a rectangular *T* × *LC* matrix with *T* ≫ *LC*,

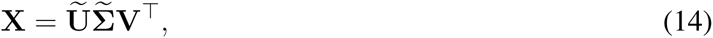

where 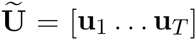 is the matrix of *output eigenvectors* (e.g. hand velocity), **V** = [**v**_1_ *…* **v**_*LC*_] is the matrix of *neural eigenvectors*, and

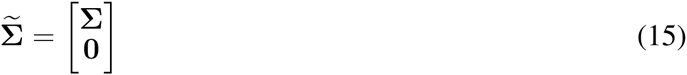

with a diagonal matrix **Σ** that has the singular values {*σ*_*i*_: *i* = 1,…, *LC*} on the diagonal. The eigenvectors {**v**_*i*_: *i* = 1,…, *LC*} form an *orthonormal basis* for the *LC*-dimensional space of possible linear decoder filters, and the eignevectors {**u**_*i*_: *i* = 1,…, *T*} form an *orthonormal basis* for the *T*-dimensional space of possible decoder outputs from decoding the neural data matrix **X** (**Hansen**, 1998; **Shlens**, 2014). The SVD formula can also be written in a ‘scalar’ form (**Shlens**, 2014)

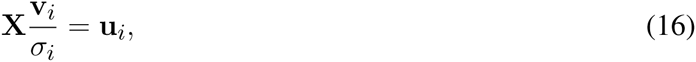

for *i* = 1,…, *LC*. Comparing this form to Equation (2), we see that indeed the vectors **v**_*i*_*/σ*_*i*_ are *candidate* linear decoder filter coefficients *tailored* for the neural data matrix **X**, with the corresponding model output represented by the vectors **u**_*i*_. To further emphasize this, we consider the Principal Component Analysis (PCA) of the neural data matrix which can be obtained – up to a scaling factor (**Shlens**, 2014) – by the eigenmode analysis of the matrix

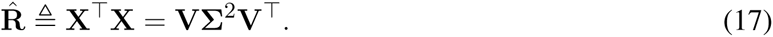

This shows that the PCs of the neural data matrix **X** are indeed the same as the eigenvectors **v**_*i*_. In other words, an *eigenmode analysis* of the neural data reveals that the candidate vectors for the filter coefficients are exactly the *eigenmodes* of the neural data matrix. Additionally, if we consider only a subset of the output eigenvectors {**u**_*i*_: *i* = 1,…, *LC*} and stack them in a matrix **U** = [**u**_1_ **u**_2_ *…* **u**_*LC*_], then the pseudo-inverse solution from Equation (3) has an equivalent form

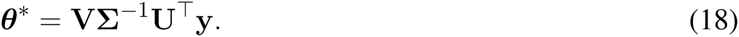

This last result can also be written in a ‘scalar’ form

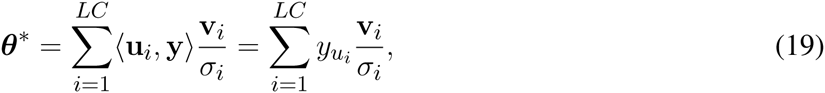

where 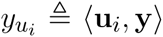 is the dot product (or degree of similarity) between the actual kinematic samples **y** and a particular decode *eigenmode* **u**_*i*_. Thus, the pseudo-inverse solution is indeed a linear combination of the neural data eigenmodes scaled by a *gain* term, 1/*σ*_*i*_.

It is worth noting that, whereas the matrix 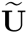 is comprised of *T* different vectors that can express *any* possible decode samples of length *T*, the matrix **U** is only comprised of *LC* vectors that can express a *subset* of such possible decode samples, with *LC* ≪ *T*. This subset is a much limited subset, comprised mainly of linear combinations of the output eignvectors **u**_*i*_ that correspond to the scaled neural eigenmodes, implying that the set of signals that can be linearly decoded from the neural data matrix **X** is a very small set as compared to the huge space of *T*-dimensional signals.

### 2.4 RIDGE REGRESSION AND REGULARIZATION

In inverse problems, the gain term 1/*σ*_*i*_ associated with the pseudo-inverse solution can become arbitrarily large whenever the singular values *σ*_*i*_ become arbitrarily small. In particular, the minimization problem (4) for very small *σ*_*i*_’s – becomes an ill-conditioned problem^4^ and the solution becomes highly sensitive to small noise in the actual data **y** (**Hansen**, 1998; **Pillonetto et al.**, 2014). Intuitively, any small degree of similarity between the measurements **y** and a particular decode eigenmode **u**_*i*_ results in an arbitrarily small dot product value *y*_*u*_*i* that is close to – but not equal to – zero. This small dot product gets sub-sequently ‘amplified’ by the gain term 1*/σ*_*i*_ and results in a large weight in the linear combination (19). This effect is particularly evident whenever the neural data record is of very short length. In such cases, the neural data matrix **X** becomes ill-conditioned, and its low-rank singular values become numerically close to zero (**Hansen**, 1998; **Bishop**, 2006; **Strang**, 2009). Moreover, such low-rank singular values typically correspond to ‘noisy’ eigenmodes of the neural data, resulting in noisy decoder filter structures and correspondingly noisy decode.

These numerical issues were recognized and addressed starting with the seminal works of **Phillips** (1962) and **Tikhonov and Arsenin** (1977) using *regularization* techniques – which are a special case of kernel-based regularization as presented in the next subsection. One of the simplest regularization techniques is the ‘Truncated SVD’ which limits the pseudo-inverse solution to the the first *M* eigenmodes (**Hansen**, 1987).

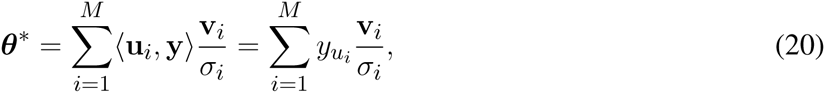

where *M* < *LC* is determined numerically. Another popular regularization technique that has been used in decoder design (**Suminski et al.**, 2010; **Collinger et al.**, 2013; **Wodlinger et al.**, 2015) is Tikhonov regularization (**Tikhonov and Arsenin**, 1977), which is also known as *ridge regression* (**Bishop**, 2006). The minimization problem (4), which is originally ill-posed, is slightly modified to become

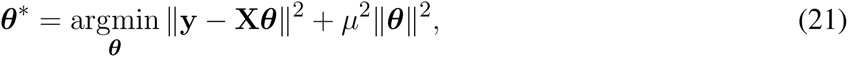

where *μ*^2^ is a regularization parameter. This regularization technique adds a ‘penalty’ term to the cost function that penalizes the magnitude of the decoder filter coefficients. The regularized solution is (**Bishop**, 2006)

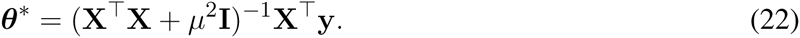

To gain more insight into the underpinnings of ridge regression, we look at the SVD of the correlation matrix **X**^T^**X** (17). Noting that *μ*^2^**I** = **V**(*μ*^2^**I**)**V**^T^, we get

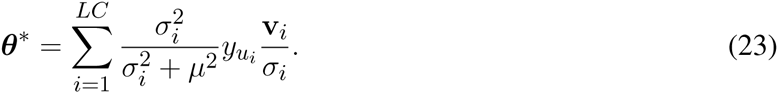

This solution converges to the pseudo-inverse solution as *μ* → 0, and converges to the zero solution (***θ*** = **0**) as *μ* → ∞, making *μ*^2^ a ‘tuning’ parameter for this solution.

As noted by **Hansen** (1998), regularization techniques are rooted in the idea of filtering out (i.e. suppressing) ‘noisy’ SVD eigenmodes. Both the truncated SVD solution and the ridge regression solution can be viewed from this ‘filtering’ standpoint. Whereas the pseudo-inverse solution has a ‘gain’ term of 1/*σ*_*i*_ for each eigenmode, the truncated SVD pre-multiplies that gain term by a filter factor

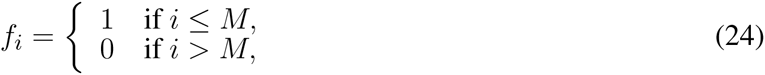

which is similar to an ‘ideal low-pass’ filter. Similarly, ridge regression uses a filter factor

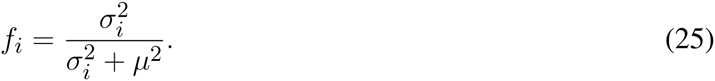

Therefore, the essence of regularization techniques is to limit the solution to a particular *subspace* with desirable properties that reflect prior knowledge. The techniques presented in this subsection limit the solution to a SVD-based subspace. This idea is generalized in the next subsection to arbitrary subspaces using kernel methods.

### 2.5 KERNEL METHODS FOR SYSTEM IDENTIFICATION

In this subsection, we present a unified framework for linear extraction of motor control signals from neural data for both biomimetic and non-biomimetic approaches. We generalize the single-input single-output (SISO) system identification methods reviewed by **Pillonetto et al.** (2014) to handle the multiple-input single-output (MISO) case^5^. This generalization is needed in order to be able to extract motor control signals by linear filtering of multiple signal sources, e.g. multiple units. We also note that this approach can be generalized to any neural signal modality, e.g. local field potentials. We further extend this framework for non-biomimetic decoding in subsection 2.10.

The first key step to this generalization is to note that the ultimate goal of the system identification procedure is to estimate the unit filter coefficients as a function of time. In other words, these unit filter coefficients are treated as functions of time that take a discrete time index as an input and produces a a unit filter coefficient as an output. Whereas in the SISO case (**Pillonetto et al.**, 2014) only *one* filter function was estimated, the MISO case involves *joint* estimation of multiple unit filter functions. This joint estimation problem is intimately related to the so-called *multi-task learning* (see **Álvarez et al.** (2012) for a review), for which two mathematically equivalent formulations exist. The first formulation is to stack all the unit filter functions into one vector-valued function and then use vector-valued kernels for the estimation, and the second – somewhat simpler – formulation is to *augment* the input space by another unit index and then use scalar-valued kernels on this new input space (**Evgeniou et al.**, 2005). For simplicity of the presentation, we use this second formulation here. Let *𝒵* = *{t*_1_, *t*_2_, *…, t*_*L*_*}* be the set of time indices for all unit filter functions *g*_*m*_(*t*). For example, *t*_1_ = 0, *t*_2_ = 1, *…, t*_*L*_ = *L -* 1 for *L* filter coefficients at *L* time indices. Let *U* = {1, 2,…, *C*} be the set of indices that index all units. Let the time-unit index tuple 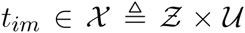 be a pair of indexes 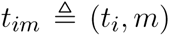 that can index all possible time indices for all unit filter functions. Then, given the input-output data, we seek to estimate a scalar function *g*: *𝓍 →* ℝ that captures all unit filter coefficients at all possible time indices such that *g*_*m*_(*t*_*i*_) = *g*(*t*_*im*_). We define a positive semidefinite *kernel*, *K*: *𝓍* × *𝓍* → ℝ

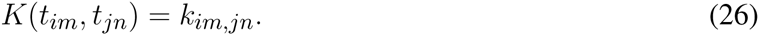

Intuitively, the scalar *k*_*im,jn*_ defines an inner product (a generalization of a vector dot product) which denotes the degree of coupling (or covariance, see next subsection) between two filter coefficients *g*_*m*_(*t*_*i*_) and *g*_*n*_(*t*_*j*_). A well-known theorem in kernel methods is the Moore-Aronszajn theorem (**Aronszajn**, 1950) which gives a one-to-one correspondence between Reproducing Kernel Hilbert Spaces (RKHS) of functions and positive semi-definite kernels. (For a lengthier treatment of RKHSs, see (**Rasmussen and Williams**, 2005; **van der Vaart et al.**, 2008; **Paiva et al.**, 2010; **Park et al.**, 2013; **Pillonetto et al.**, 2014).) For our purpose, the implication of this theorem is that, once a kernel is defined, the space of possible unit filter functions *g*_*m*_(*t*) is restricted to the corresponding RKHS *ℋ*. Since this space encodes some notion of ‘smoothness’ of its member functions (**Rasmussen and Williams**, 2005; **Pillonetto et al.**, 2014), we immediately see the practical advantage of such restriction: the unit filter functions are restricted to ‘smooth’ functions of time. In essence, the space *ℋ* corresponding to the defined kernel becomes the *hypothesis space* within which we seek unit filter functions that minimize the error between the decode and the actual kinematics. Adopting a RKHS as a hypothesis space takes care of the ill-conditioning of the minimization problem at hand, since the modified minimization problem (stated below) is well-conditioned (**Pillonetto et al.**, 2014).

Referring to Figure 1, we define a unit output *z*_*m*_(*t*) as the output of filtering the *m*^th^ unit binned spike counts^6^, *x*_*m*_, with the corresponding unit filter *g*_*m*_. Using the ‘functional’ notation as in **Pillonetto et al.** (2014), a unit filter output is written as

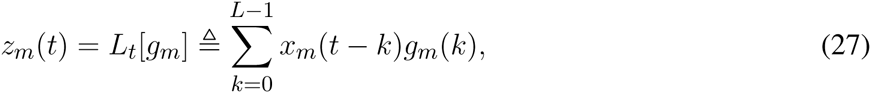

where *L*_*t*_[*g*_*m*_] is a ‘functional’ (i.e. a function that takes another function as an input) that represents a unit output at time *t*. With this notation, the decode is expressed as

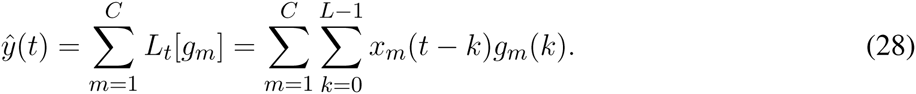

This equation – with minor notational changes – is the same as the optimal linear decoder presented earlier in (1). With this notation, the variational minimization problem for the MISO case can be succinctly written as

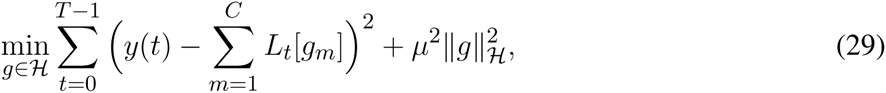

where 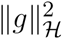 is the induced norm of the RKHS *ℋ* (**Pillonetto et al.**, 2014).

This abstraction is quite powerful. It can be readily extended to filters of infinite duration or even continuous time filters. Nonetheless, we greatly simplify the presentation here to give the algorithmic methods and refer the interested reader to **Pillonetto et al.** (2014) and **Appendix 1** for more details on the case of infinite-duration filters. Corresponding to the kernel *K*, we can construct a *kernel matrix* **Q** (**Bishop**, 2006; **Pillonetto et al.**, 2014) at a finite set of time-unit indices. The construction of the kernel matrix is similar to the construction of the correlation matrix **R** (see subsection 2.1), by stacking *C* × *C* submatrices, **q**_*mn*_, into a big matrix **Q**. A submatrix **q**_*mn*_ is a *L* × *L* covariance matrix between filter coefficients of units *m* and *n* at *L* different time indices (from the set *Z*). This matrix **Q** can be interpreted as a covariance matrix as presented in the next subsection. The individual elements of a submatrix **q**_*mn*_ are given by

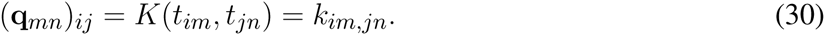

The modified minimization problem can be simplified – with a slight abuse of notation – by stacking the coefficients of all the finite-duration unit filters in one vector ***θ*** as in (2)

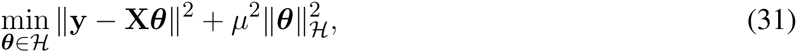

where the notation ***θ*** ∈ *ℋ* means that the elements of ***θ*** are taken as samples from the scalar function *g ∈ H*, and 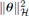 is simply used to denote the norm 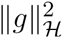. This latter norm is greatly simplified (**Pillonetto et al.**, 2014) for finite-duration filter functions as^7^

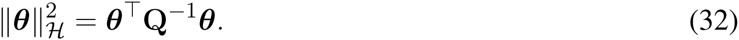

Therefore, this squared norm is indeed the squared Mahalanobis distance between the vector ***θ*** and a multivariate Gaussian distribution with zero-mean and a covariance matrix **Q** (**Rasmussen and Williams**, 2005). By comparing this norm to the corresponding *ℓ*_2_-norm employed in ridge regression (Equation (21)), we see that both ridge regression and kernel-based solutions employ a penalty term. Whereas ridge regression penalizes large decoder coefficients uniformly across all eigenmodes, kernel-based regularization only penalizes large decoder coefficients in the directions of eigenmodes with small singular values (See **Remark 1** in **Pillonetto et al.** (2014) for SVD interpretation of a kernel matrix).

**Table 1.**
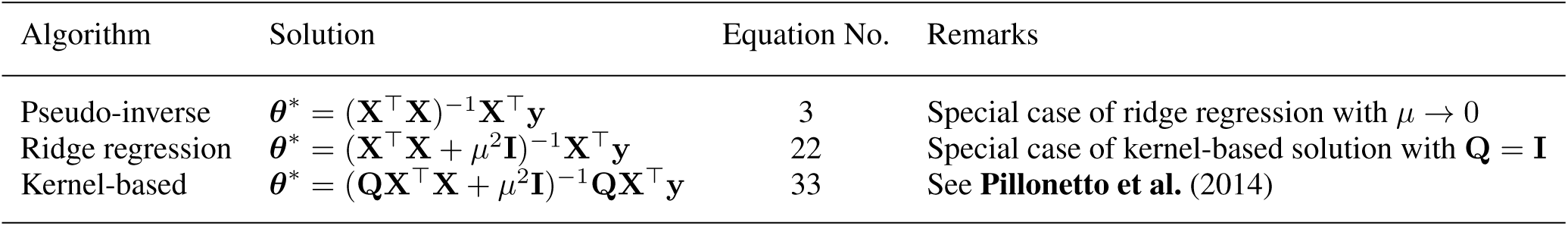
Summary of the optimal linear decoding algorithms

The solution to this minimization problem (see **Appendix 2** in the supplementary material) is

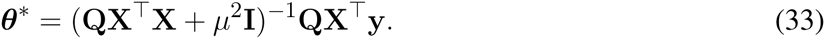

We conclude this subsection by a few remarks on the connection between the pseudo-inverse, ridge regression, and kernel-based solutions. The ridge regression solution is a special case of the kernel-based solution with **Q** = **I**. In other words, the ridge regression solution assumes no coupling between unit filter coefficients across different time indices, and no coupling between the different unit filters across units. Further, the pseudo-inverse solution is a special case of the ridge regression solution with *μ* → 0. These remarks together with an overview of these solutions are summarized in Table 1.

### 2.6 BAYESIAN INTERPRETATION

Both ridge regression and kernel-based solutions have an equivalent probabilistic interpretation in a Bayesian framework (**Rasmussen and Williams**, 2005; **Bishop**, 2006; **van der Vaart et al.**, 2008; **Pillonetto et al.**, 2014). Consider the generative data model **y** = **X *θ*** + ***ϵ***. In a Bayesian treatment, the parameters to be estimated – which are the decoder filter coefficients – are treated as random variables. In particular, if we assume for the random vector ***θ*** a Gaussian distribution with zero mean and covariance matrix **Q** – same as the kernel matrix – then the combined data and parameters vector has a joint-Gaussian distribution (**Pillonetto et al.**, 2014)

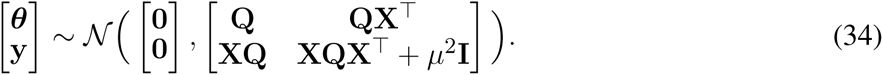

The kernel matrix **Q** acts as the covariance matrix of a *prior* distribution on the parameters. Moreover, standard results from Gaussian-process regression directly give us a *posterior* distribution on the parameters after observing the data **y**

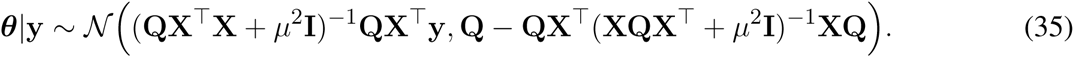

Thus, the *mean* of the posterior distribution – which is also the mode or the maximizer of the posterior in the Gaussian case – is precisely the kernel-based solution (33). Moreover, the Bayesian treatment gives us uncertainty bounds that can be calculated from the posterior covariance. Thereby, on the one hand, the pseudo-inverse solution can be interpreted as the *maximum likelihood* (ML) solution to the parameter estimation problem. On the other hand, the kernel-based solution – as well as the ridge regression solution as a special case – can be interpreted as the *maximum a posteriori* (MAP) solution. The quality of the solution for short data records largely depends on the chosen kernel matrix **Q**. **Pillonetto et al.** (2014) proved that the optimal kernel matrix should have exactly one eigenmode in the direction of the ‘true’ solution. Of course, since the ‘true’ solution is never known beforehand, we need to carefully design a kernel matrix that reflects prior knowledge about how the solution *should look like* in terms of its eigenmode directions, and the extent of how large the filter coefficients should be in these particular directions.

### 2.7 PROPOSED KERNELS FOR BIOMIMETIC DECODING

As we noted earlier, ridge regression is equivalent to an identity kernel matrix **Q** = **I**. However, this kernel structure assumes that all decoder filter coefficients are *uncorrelated*. The hypothesis that we put forward is that optimal unit filter functions possess a well-defined structure that coincides with the eigenmodes of the neural data (**Badreldin et al.**, 2013). This hypothesis is motivated by several empirical observations and theoretical considerations. First, from a *matched filter* perspective, we note that unit filter functions operate as ‘correlators’ that look for specific neural patterns of co-activation in the input binned spike counts. Second, for the case of decoding natural velocity trajectories, we conjecture that the characteristic velocity peaks with smooth profiles that are typically observed in primate reaching behavior can be decoded using ‘matched filters’ that operate on the neural input (**Badreldin and Oweiss**, 2014). Third, regularized decoders calibrated using biomimetic data typically possess characteristic unit filter structures with exponentially decaying oscillations. This empirical observation is also reported elsewhere (**Fagg et al.**, 2009; **Willett et al.**, 2012; **Flint et al.**, 2012). Fourth, single-unit dynamics quantified by peri-event time histograms (PETHs) constructed around the velocity peaks – as shown in the results – exhibit characteristic structures that resemble the structures of unit filters. Fifth, as was shown in subsections 2.3 and 2.4, unit filter functions are markedly expressed as a linear combination of neural co-activation eigenmodes.

Based on this hypothesis, the first kernel matrix we propose is to use the estimated unnormalized neural covariance^8^ matrix 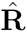 as a kernel matrix. This gives more preference to filter functions that mimic neural eigenmodes. However, this also gives higher weights to units that exhibit strong modulations from their baseline firing rates, possibly masking other units that are naturally more silent or do not modulate as strongly. This potential drawback motivates the second kernel matrix that we propose, which is a diagonal-normalized version of the first one. By normalizing the kernel matrix such that all diagonal entries are ones, this kernel gives more uniform weights to all units. The diagonal-normalized kernel matrix is calculated from the matrix 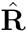 in two steps. First, a symmetric matrix is constructed using the reciprocal of the diagonal elements of the matrix 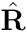

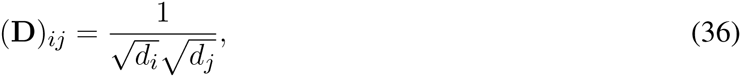

where {*d*_*i*_: *i* = 1, 2,…, *LC*} are the diagonal elements of the matrix 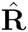. Second, the diagonal-normalized kernel matrix, 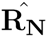, is calculated using element-wise matrix product (or Hadamard product) of 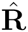 and **D**.

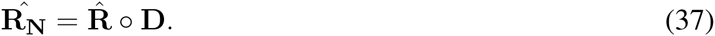

Finally, for numerical conditioning, any diagonal element in 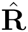 that is strictly less than one is replaced by one in the matrix **D**.

At this point, it is informative to examine the SVD of the kernel-based solution in (33). With the first proposed kernel, **Q** = 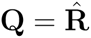, we get

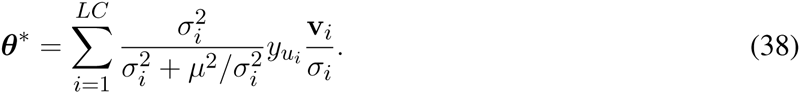

Hence, the SVD filter factor of this kernel matrix is

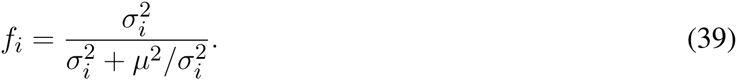

**Table 2.** Summary of the SVD formulas of different decoding algorithms

| Algorithm | Full Solution | Equation No. | Truncated Solution |
| --- | --- | --- | --- |
| Pseudo-inverse | $\theta^* = \sum_{i=1}^{LC} y_{u_i} \frac{\mathbf{v}_i}{\sigma_i}$ | 19 | $\theta^* = \sum_{i=1}^M y_{u_i} \frac{\mathbf{v}_i}{\sigma_i}$ |
| Ridge regression | $\theta^* = \sum_{i=1}^{LC} \frac{\sigma_i^2}{\sigma_i^2 + \mu^2} y_{u_i} \frac{\mathbf{v}_i}{\sigma_i}$ | 23 | $\theta^* = \sum_{i=1}^M \frac{\sigma_i^2}{\sigma_i^2 + \mu^2} y_{u_i} \frac{\mathbf{v}_i}{\sigma_i}$ |
| Neural covariance kernel | $\theta^* = \sum_{i=1}^{LC} \frac{\sigma_i^2}{\sigma_i^2 + \mu^2 / \sigma_i^2} y_{u_i} \frac{\mathbf{v}_i}{\sigma_i}$ | 38 | $\theta^* = \sum_{i=1}^M \frac{\sigma_i^2}{\sigma_i^2 + \mu^2 / \sigma_i^2} y_{u_i} \frac{\mathbf{v}_i}{\sigma_i}$ |

Compared to the ridge regression filter factor in Equation (25), we see that the weight of the regularization parameter is varying as a function of the rank of the singular values. High-ranked singular values, which are typically large, dilute the regularization parameter, thereby giving more SVD gain to the corresponding eigenmodes, whereas this effect is reversed for low-ranked singular values which are typically small. We conclude by summarizing the SVD formulas of different truncated and non-truncated (i.e. full) solutions to the decoder design problem as presented in Table 2.

### 2.8 UNIT-FILTER MATCH METRICS

From a ‘matched filter’ standpoint, we hypothesize that the structure of the single-unit firing rates around the natural velocity peaks should match the structure of the corresponding unit filter functions. We propose to quantify this ‘degree of matching’ using two different metrics. By constructing single-unit PETHs in one-second windows prior to the times of the velocity peaks with the same bin size used for the unit filters, we can compute Pearson correlation coefficients *r*_*m*_ between single-unit PETHs and corresponding unit filter functions. The first metric we propose is

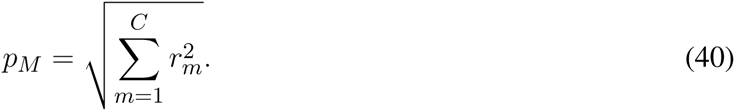

We refer to this metric as the unit-filter match *magnitude*. The second metric – which relies on the statistical significance of these correlations – is the fraction of units for which *r*_*m*_ is significantly different than zero (*p* < 0.05) under a *t*-test. We refer to this metric as the unit-filter match *unit fraction*.

### 2.9 PERFORMANCE METRICS OF DECODERS

Performance of different decoders has been traditionally assessed based on the ability of a decoder model to calculate a decode that is as close as possible to actual test data. For this performance aspect, we use the *R*^2^ metric – which is known in the literature as the *coefficient of determination* and also as the *fraction of variance accounted for* (**Fagg et al.**, 2009)

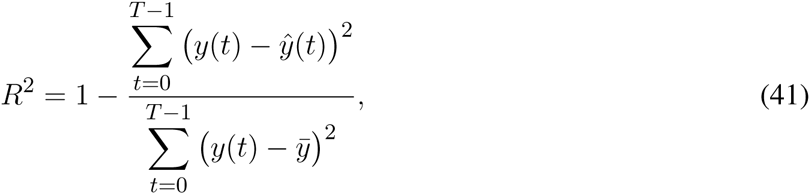

where 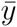 is the mean of the actual data samples, *y*(*t*). This metric is a unitless metric in the range [–∞, 1], with the upper limit indicating perfect model output.

In addition to this standard metric, we propose five additional metrics that can be used to judge the online performance of a decoder and its fitness for real-time robotic control – even before closing the loop with this decoder. In other words, we conjecture that deterioration in performance as assessed by these metrics can inform a decoder re-calibration decision. The first metric we propose is to quantify the *signal-to-noise ratio* (SNR) of a kinematic signal. In particular, we note that hand velocity trajectories during reaching by primates – which is the main motor control signal we are attempting to extract – typically exhibit characteristic peaks that are approximately bell-shaped (**Soechting**, 1984; **Flash and Hogan**, 1985). These characteristic peaks are also referred to as submovement primitives (**Gowda et al.**, in press). Aside from these characteristic peaks, a velocity trajectory mainly consists of small-amplitude oscillations. Consequently, the histogram of the absolute values of a hand velocity trajectory in one degree of freedom (DOF) is typically *bimodal*, where one mode around *zero* is due to the ‘noise’ component, and the other mode – which is typically a *heavy tail* of the histogram – is due to the characteristic hand velocity peaks, i.e. a ‘signal’ component. Therefore, discriminating the ‘signal’ component from the ‘noise’ component becomes a binary detection problem that is mathematically similar to detection of spike waveforms in extracellular potentials (**Oweiss and Aghagolzadeh**, 2010). A ‘spike’ waveform to be detected here, however, is a characteristic velocity peak – which we refer to as a ‘velocity spike’. We therefore propose to detect these velocity spikes in a similar manner to detection of extracellular spikes by simple thresholding (**Oweiss and Aghagolzadeh**, 2010; **Aghagolzadeh et al.**, 2014). For a hand velocity signal *y*(*t*) – or in general for any kinematic signal that exhibits characteristic ‘spikes’ – we detect velocity spikes as a waveform ‘snippet’ between two event times. The first (second) event time is the time at which the absolute signal |*y*(*t*)| crosses above (below) a predefined threshold. Algorithmically, the first event time is

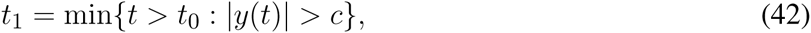

where *c* is a predefined threshold and *t*_0_ is some initial time that can be taken as the start of a record or the end time of a previously detected velocity spike. The second event time is

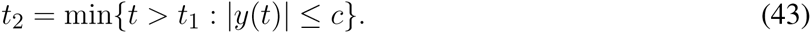

A velocity spike snippet is a collection of absolute velocity samples stacked up in a vector

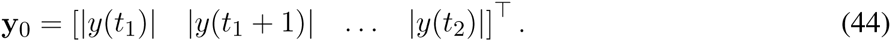

Next, we define the SNR of velocity spikes similarly to the SNR of spikes detected in extracellular potentials

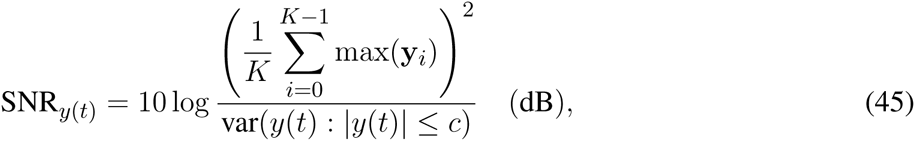

where *K* is the number of extracted velocity spikes. In other words, it is the ratio of the squared average peak of velocity spikes (‘signal’) to the variance of the subthreshold velocity values (‘noise’). The ‘noise’ threshold *c* is calculated as the *center of mass* of the empirical probability mass function (i.e. the normalized histogram) of the absolute velocity values |*y*(*t*)|. This center of mass is mathematically equivalent to an expectation taken over the empirical probability mass function of the values |*y*(*t*)|

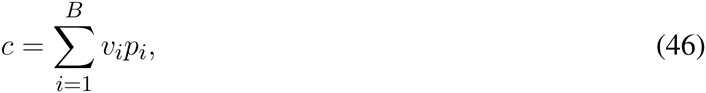

where *v*_*i*_ is a center of histogram bin number *i* and *p*_*i*_ is the empirical probability associated with this bin. Numerically, the total number of bins *B* can be taken as the ceiling integer to the square root of the number of velocity samples 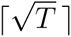 (**Maciejewski**, 2011).

The second metric we define, which has been introduced earlier (**Badreldin et al.**, 2013; **Badreldin and Oweiss**, 2014), is the average number of zero-crossings per second, *ζ*, which is calculated as

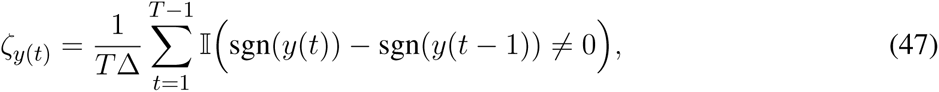

where Δ is the bin width of the kinematic signal *y*(*t*), 𝕀 (·) is the indicator function, and sgn(·) is the signum function.

The third metric, which we call the *filter latency*, quantifies the average feed-forward delay introduced by the decoder filters. It is calculated using the *phase response* of the individual unit filters of a decoder. Denote the frequency response of a unit filter function *g*_*m*_(*t*) by *G*_*m*_(*jω*), then the phase response of a unit filter is

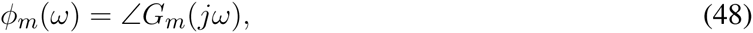

and the filter delay as a function of frequency – also known as the *group delay* of a filter (**Oppenheim and Willsky**, 1997) – is given by the negative of the first-order derivative of the phase response, 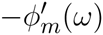 We define the filter latency as the average filter delay across all frequencies and across all units

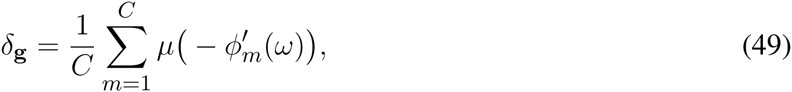

where *μ*(·) is the mean of a filter’s group delay.

The fourth metric – which was also introduced earlier in **Badreldin et al.** (2013) – quantifies the number of units that dominate the computation of the decode. Since a typical decoder does *not* assign the same weights to all units, it *may* be desirable – from a pragmatic standpoint – to have more unit contributions to the computation of the decode. This can help in maintaining a decoder fixed across multiple days – even if some of the decoder units disappear for a few days (**Heliot et al.**, 2010; **Eleryan et al.**, 2014). A unit’s contribution to the decode can be quantified in terms of its output *z*_*i*_(*t*). By stacking up all the unit output samples in one vector **z**_*i*_, we define the unit contribution index *ν* as the fraction of units whose outputs constitute more than 90% of the total sum magnitudes of all unit outputs. Algorithmically, let

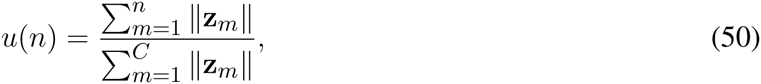

where the unit numbers, *m* = 1,…, *C*, are sorted in descending order of unit output magnitude. From this, *v* is calculated by

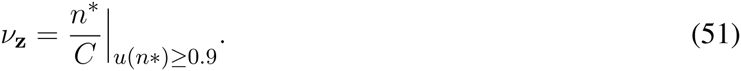

We note that this metric assesses unit contributions in a way that is comparable to the ‘neural push’ unit contribution metric proposed by **Stavisky et al.** (2015).

The fifth metric we propose is a variation on the sample skewness that we used in earlier work (**Badreldin et al.**, 2013; **Badreldin and Oweiss**, 2014). Sample skewness roughly quantifies the degree of non-symmetry of a decode distribution. A symmetric decode is desirable because it spans the entire task space with no bias to particular directions. However, the sample skewness metric is only limited to one-dimensional signals, and it has consistency problems since a sample skewness value of zero does not necessarily imply that the distribution is symmetric (**Székely and Móri**, 2001). To address these problems, we make use of another metric of asymmetry of a distribution. In particular, **Székely and Móri** (2001) proved that for two random vectors **x** and **y** that are independently sampled from the same distribution, the following result holds

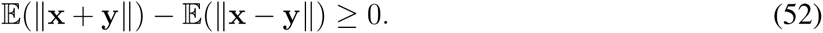

Which can be rearranged as

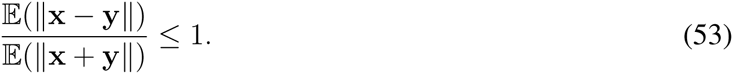

Moreover, equality holds if and only if the underlying distribution is *perfectly symmetric*, i.e. when the random variables x and –x have exactly the same distribution. Using this fact, we define a sample-based symmetry metric

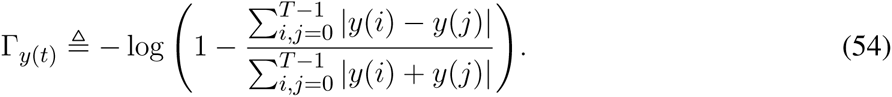

This metric is in the range [0, ∞], with the upper limit indicating a perfectly symmetric distribution around zero (**Székely and Móri**, 2001).

### 2.10 EXTENSION TO NON-BIOMIMETIC DECODING

One advantage of using kernel methods in decoder design is the possibility to exploit the same proposed kernels for design of non-biomimetic decoders in the absence of actual or observed kinematic signals (**Badreldin et al.**, 2013; **Badreldin and Oweiss**, 2014). In a biomimetic approach, the cost function is defined in terms of an error metric between the decode and the actual kinematics. However, in the absence of the latter, it is necessary to optimize a different cost function. In **Badreldin and Oweiss** (2014), we proposed to maximize the kurtosis of the decode as a proxy for its SNR. Here, we instead propose to directly maximize the SNR of the decode as defined in (45). Similar to **Badreldin and Oweiss** (2014), we propose a constrained maximization problem

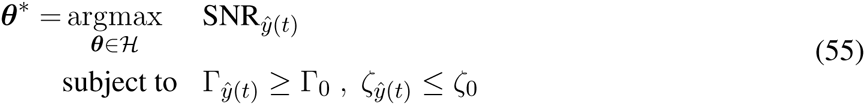

where 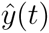 is the decode, and the parameters Γ_0_ and *ζ*_0_ are determined from prior knowledge. This constrained maximization problem attempts to maximize the SNR of the decode, 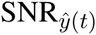, while maintaining a symmetric motor control signal, 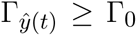, and bounding the average number of zero-crossings per second, 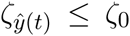. This last metric is roughly related to the amount of ‘noise’ in a signal since – in the absence of velocity spikes – the number of zero-crossings per second approximately represents the fundamental frequency of the ‘noise’ component.

The choice of a particular kernel fully defines the *hypothesis space **H*** (see subsection 2.5). In **Badreldin et al.** (2013), we made use of the neural covariance kernel, whereas in **Badreldin and Oweiss** (2014) we made use of a variation on the diagonal-normalized kernel – albeit with infinite duration, high-resolution, filters. In both cases, we found that it is computationally more feasible to use truncated kernels that are comprised of a finite number of basis functions (see Table 2). Hence, we propose to use the truncated versions of the neural covariance and the diagonal-normalized kernels presented in subsection 2.7. Since kernel methods allow the use of prior knowledge to inform how the solution should look like, we propose to choose the unit filters basis functions that produce decode components with *desirable* features for motor control. This way, we favor particular directions in the solution space that better meet the constraints in (55). Following the design methodology in **Badreldin et al.** (2013), we limit the hypothesis space to neural eigenmodes that produce decode components that satisfy the constraint 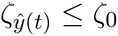.

Another important characteristic of the solution is that the unit filters as a function of time should have an envelope that decays exponentially. The importance of this envelope stems from the relative weighting it introduces with regard to immediate neural input history versus far history. Putting the emphasis of the filter weights on immediate neural input history results in decoders that have lower latencies – as demonstrated in the results. To this end, we propose to synthetically ‘taper’ the chosen neural eigenmodes by windowing their filter functions with a predefined time window function

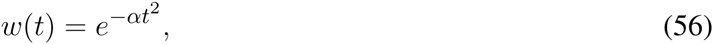

where *α* is chosen to control the decay rate. Example window functions for practical values of *α* are shown in Supplementary Figure 1. In practice, a particular value of *α* to achieve a 50% decay of the filter weights at a predefined time point *t*_0.5_ can computed as

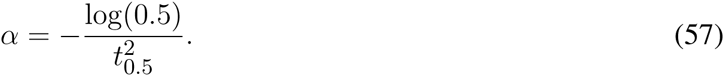

Therefore, *t*_0.5_ can be computed in such a way as to mimic the structure of regularized biomimetic decoders. To this end, it is useful to define a *decoder half-RMS point* for a particular decoder. By first stacking up all unit filter coefficients at each time point in a vector

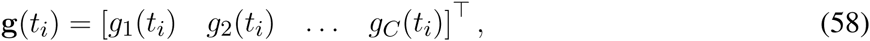

we define the decoder half-RMS point

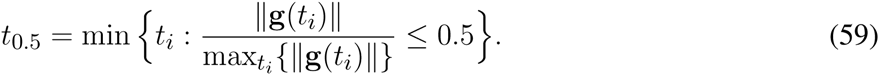

### 3 RESULTS

In this section, we demonstrate the underpinnings and performance of the proposed algorithms for both biomimetic and non-biomimetic decoding using publicly available data from the Database for Reaching Experiments and Models (DREAM) (**Walker and Kording**, 2013), hosted by the Collaborative Research in Computational Neuroscience (CRCNS) website (**Teeters et al.**, 2008). In particular, we use synchronized kinematic and neural data recorded during an eight-target center-out task from one rhesus macaque (monkey C in **Flint et al.** (2012)). The monkey was trained to perform a center-out reaching task while grasping a two-link manipulandum. The monkey was required to reach to one of eight 2-cm^2^ targets spaced at 45° intervals around a circle of radius 10 cm. Neural data were recorded using a 96-channel microelectrode array implanted in M1. More details on the experimental procedure and the collection of data can be found in **Flint et al.** (2012).

#### 3.1 KERNELS AND REGULARIZATION

As noted in subsection 2.6, kernel methods are equivalent to a Bayesian approach where a prior distribution on the decoder filters is combined with the likelihood function from the data to get a posterior distribution, from which the decoder filters are obtained as the distribution mean. Here we demonstrate the properties of the decoder filters sampled from different priors corresponding to the proposed kernels in order to justify the choice of these kernels.

Figure 2 demonstrates the spatiotemporal structure of random samples taken from different kernels. The first kernel is the identity matrix, which is the kernel used in ridge regression (RIDG). The second kernel is the truncated neural covariance kernel (COV|T). The third kernel is the diagonal-normalized, truncated neural covariance kernel (COV|N|T). The fourth kernel is the windowed, truncated neural covariance kernel (COV|T|W). The fifth kernel is the diagonal-normalized, windowed, truncated neural covariance kernel (COV|N|T|W). All truncated kernels in this figure are constructed from the first 13 basis functions to emphasize slow dynamics. The tapered kernels used a time window function with *t*_0.5_ = 0.35 (sec). The ‘typical’ samples demonstrated in panel **B** are selected from 100 samples taken from the respective distributions, where the selected sample has the minimum Mahalanobis distance to the respective distribution. The spatial structure is captured in terms of the unit filter magnitude (panel **C**), and the temporal structure is captured in terms of the filters RMS as a function of time (panel **D**).

**Figure 2.**
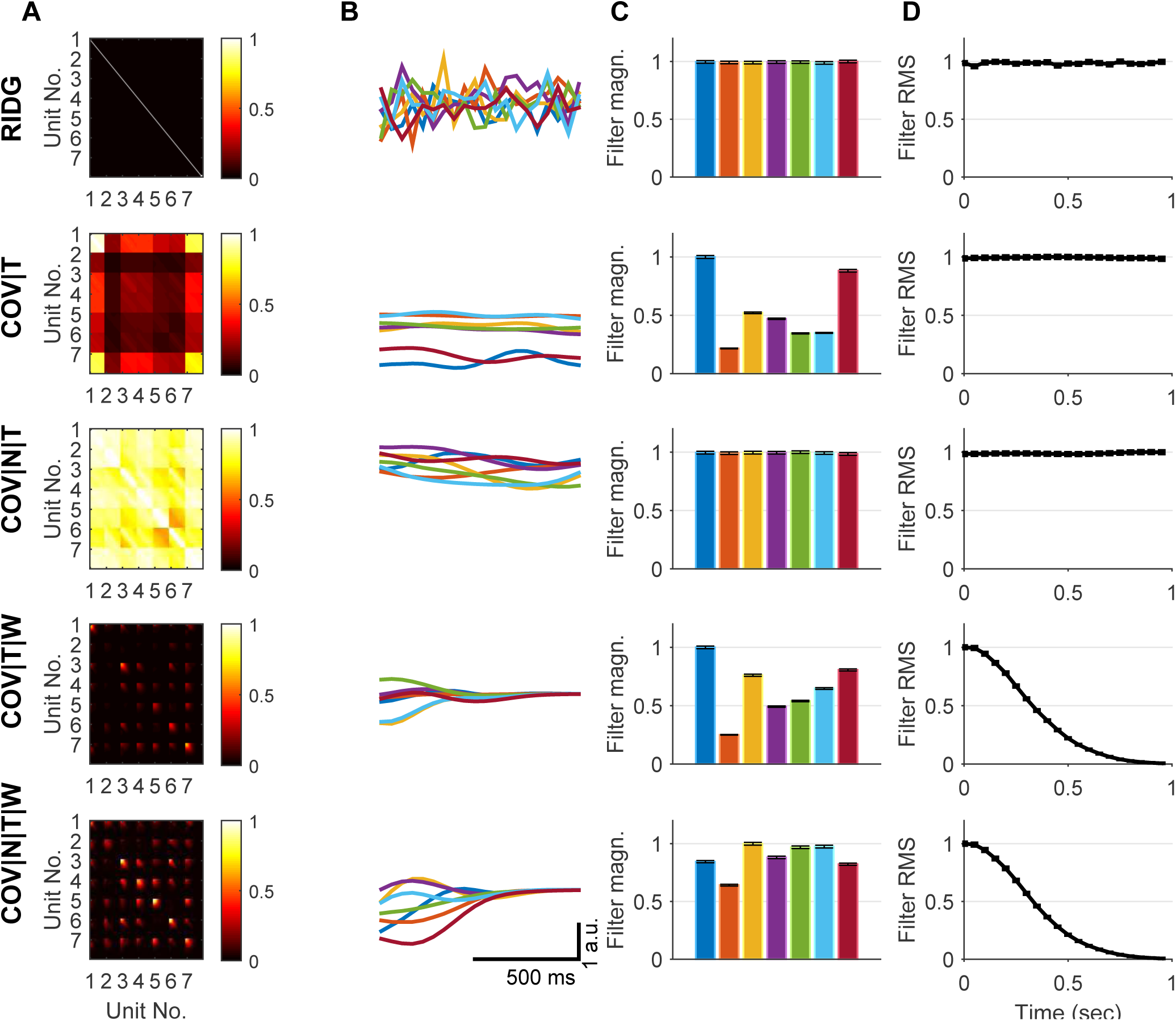
Sampling from prior distributions. (**A**) Visualization of the absolute values of the kernels as heat-maps. (All kernel matrices were normalized to have a maximum absolute value of one.) (**B**) A ‘typical’ sample from the corresponding prior distribution with zero mean and covariance equal to the kernel matrix in **A**. (Colors represent different units.) (**C**) Unit filter magnitude for each unit, averaged over 1000 samples, and normalized to have a maximum of one. (Error bars represent standard error of the mean.) (**D**) Filter RMS values as a function of time, averaged over 1000 samples, and normalized to have a maximum of one. (Error bars represent standard error of the mean.)

The temporal structure of the RIDG kernel samples is similar to that of white noise. This is because the off-diagonal entries of the kernel are all zeros which represents uncorrelated filter coefficients. Moreover, samples from this kernel have unit filters that have the same magnitude (on average), because the kernel diagonal entries are all ones. Lastly, the average temporal profile of the kernel samples is flat. The COV|T kernel samples typically have non-uniform unit filter magnitudes, where some unit filters have higher magnitudes than others. This effect can be seen in Figure 2 (**B** and **C**). The reason is that the unnormalized neural covariance kernel assigns relative ‘variance’ values for the unit filters that are similar to the relative variances of the unit firing rates. Samples from this kernel clearly show structured temporal dynamics as seen in Figure 2**B**. However, this effect is not clear in Figure 2**D**. The reason is that, although this kernel encodes temporal covariations between different unit (relative phase coupling), it does not encode absolute phase values. Hence, samples taken from this kernel typically ‘peak’ at relatively fixed locations, but not absolutely fixed locations. The COV|N|T kernel possesses similar properties, except that it assigns more uniform filter magnitudes to all units. The ‘tapered’ kernels COV|T|W and COV|N|T|W are similar to their respective untapered versions, except that the temporal profiles of the unit filters are forced to follow the time window function as revealed in Figure 2**D**.

Figure 3 illustrates a different aspect of kernel methods based on SVD analysis. The first few (low-ranked) neural eigenmodes typically consist of slow oscillations, whereas the last (high-ranked) neural eigenmodes are high-frequency oscillations and ‘noise’. By comparing pseudo-inverse (PINV), RIDG, and COV solutions in terms of their SVD gain and filter factors, we note that they give different weights to different neural eigenmodes. Whereas PINV solution amplifies high-ranked neural eigenmodes – giving rise to ‘noisy’ filters, RIDG and COV solutions attenuate such high-ranked eigenmodes – giving rise to less noisy filters.

**Figure 3.**
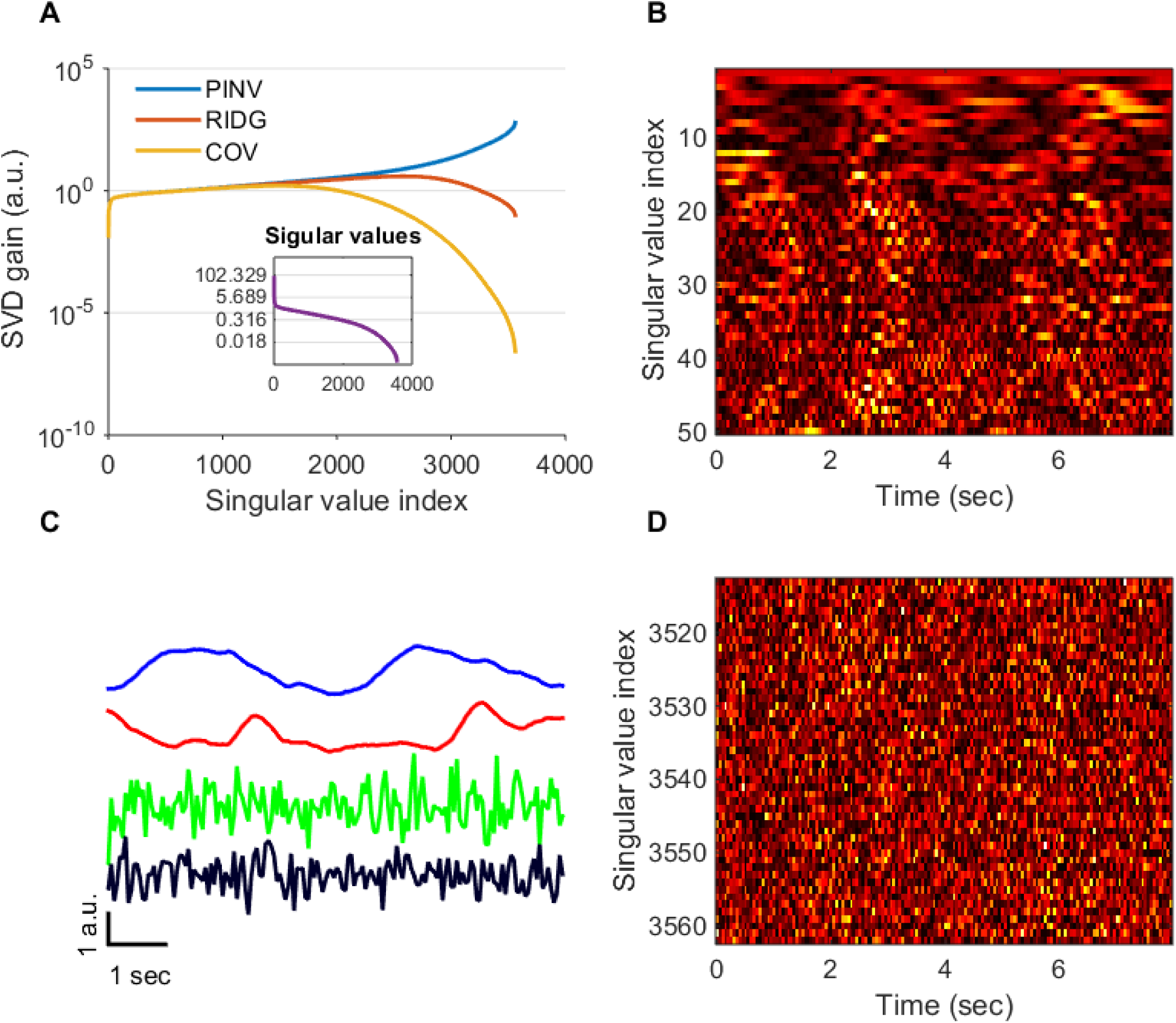
Neural eigenmodes and the effect of regularization. (**A**) The gain term given by the different algorithms to the neural eigenmodes (SVD basis vectors). Whereas the pseudo-inverse solution gives large weights to lower-ranked SVD basis, the regularized solutions give less weight to such indices which typically correspond to high-frequency output eigenmodes. Truncated versions of these algorithms follow the same pattern up to the point of the truncation index, after which the gain drops to zero. Inset shows a typical distribution of the singular values that give rise to such gain terms. Low-ranked singular values are typically very small, specially for dominantly low-frequency inputs. Such small singular values, when inverted, give rise to large SVD gains as seen in the main plot. (**B**) and (**D**) illustrate a typical structure of the top-50 (**B**) and bottom-50 (**D**) output eigenmodes. (**C**) The first two output eigenmodes from **B** and the last two output eigenmodes from **D**. These plots demonstrate the ubiquitous empirical observation that, for some bandlimited low-frequency input, the low-ranked SVD output eigenmodes are dominated by low-frequency components whereas the high-ranked ones are dominated by high-frequency components.

#### 3.2 BIOMIMETIC DECODING PERFORMANCE

We use all the data from **Flint et al.** (2012) to evaluate the offline decoding performance of natural movement from spike data. The data set comprises four files, where each file contains one or more sessions (approximately 10 minutes each) of the center-out task, totaling 11 sessions. Each session consists of natural reach ‘trials’. For each session, we concatenated all trials back to back and resampled the kinematic data on a uniform time grid of 50-millisecond time bin width. We used the spike counts with the same bin width as the kinematics to decode the first component of the natural hand velocity using one-second-long (20 time bins) decoder filters. We used all data from all 11 sessions, without excluding any data segments. We employed a variation on the generalized cross-validation scheme (**Bishop**, 2006) for reporting decoding performance. Each session was divided into 10 ‘blocks’ of data, where each block is approximately one minute. We used three blocks for ‘training’ the decoders using all algorithms, i.e. to estimate the decoder filters. Then, we used seven blocks as ‘test’ data to report decoding performance. Each session had 10 different training/test arrangements as illustrated in Supplementary Figure 2, resulting in a total of 110 block arrangements on which we report performance. We further divided the three blocks of training data into two blocks for estimating the neural and kinematic covariance matrices, 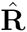 and 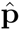, and one block for tuning the regularization parameters. We compare the three main algorithms from Table 1, namely: PINV, RIDG, COV, and COV|N. Additionally, we also compare the ‘truncated’ versions of these algorithms by using only the first *M* basis vectors from the SVD analysis. We refer to these truncated version using the same mnemonic with added ‘|T’ suffix, i.e. PINV|T, RIDG|T, COV|T, and COV|N|T, respectively. Finally, since PINV does not have any regularization parameters, its best performance is achieved with longer data records. Therefore, for PINV, we used the entire three blocks of training data with no further partitioning.

Supplementary Figure 3 demonstrates – on a typical block arrangement – the procedure for tuning the regularization parameters for all algorithms (except for PINV which does not have a regularization parameter). The non-truncated algorithms have only one regularization parameter, *μ*^2^. The value of this regularization parameter is chosen to maximize generalization performance (in terms of *R*^2^) on unseen data. For the truncated algorithms, two parameters need to be tuned, which are the truncation order *M* and the regularization parameter *μ*^2^. We select these parameters using a (possibly suboptimal) greedy algorithm – by first selecting *M* followed by *μ*^2^.

Using these block arrangements and parameter tuning procedures, we conducted a multi-faceted analysis of the decoding performance of all algorithms. Performance data from all 110 block arrangements are presented in Figure 4. We emphasize that optimization tuning of all algorithms was done to maximize only one metric, which is the *R*^2^. Variations in performance in all other metrics are mainly reflecting the different priors. We used Wilcoxon signed rank test with Bonferroni correction for multiple comparisons for conservative post-hoc comparison of algorithmic performance. On the one hand, PINV performed significantly worse than all others in terms of *R*^2^ with its median near zero (*p* < 0.05). Moreover, it had significantly worse decode SNR than all others (*p* < 0.05) with a median of 5.38 dB. Similarly, its decode had significantly higher zero-crossings per second (*p* < 0.05) with a median of 3.77 zero-crossings/sec. Additionally, the average decoder filter latency for PINV decoders was significantly worse than all others (*p <* 0.05) with a median of 0.43 sec. Finally, PINV decoders had significantly the highest unit contribution index among all others (*p* < 0.05) with a median of 0.69. On the other hand, COV|N demonstrated the best overall performance – it outperformed all other algorithms across all block arrangement in terms of *R*^2^ (69%), SNR (37%), zero-crossings per second (47%), and low latency (58%). COV|N performed significantly better than all others in terms of *R*^2^ (*p* < 0.05) with a median of 0.68, decode SNR (*p* < 0.05) with a median of 6.12 dB (except for COV|N|T not significant), zero-crossings per second (*p* < 0.05) with a median of 2.06 sec^*-*1^, and low latency (*p* < 0.05) with a median of 0.29 sec. COV|N had a unit contribution index that is significantly higher (*p* < 0.05) than all others except for PINV and COV|N|T with a median of 0.44.

**Figure 4.**
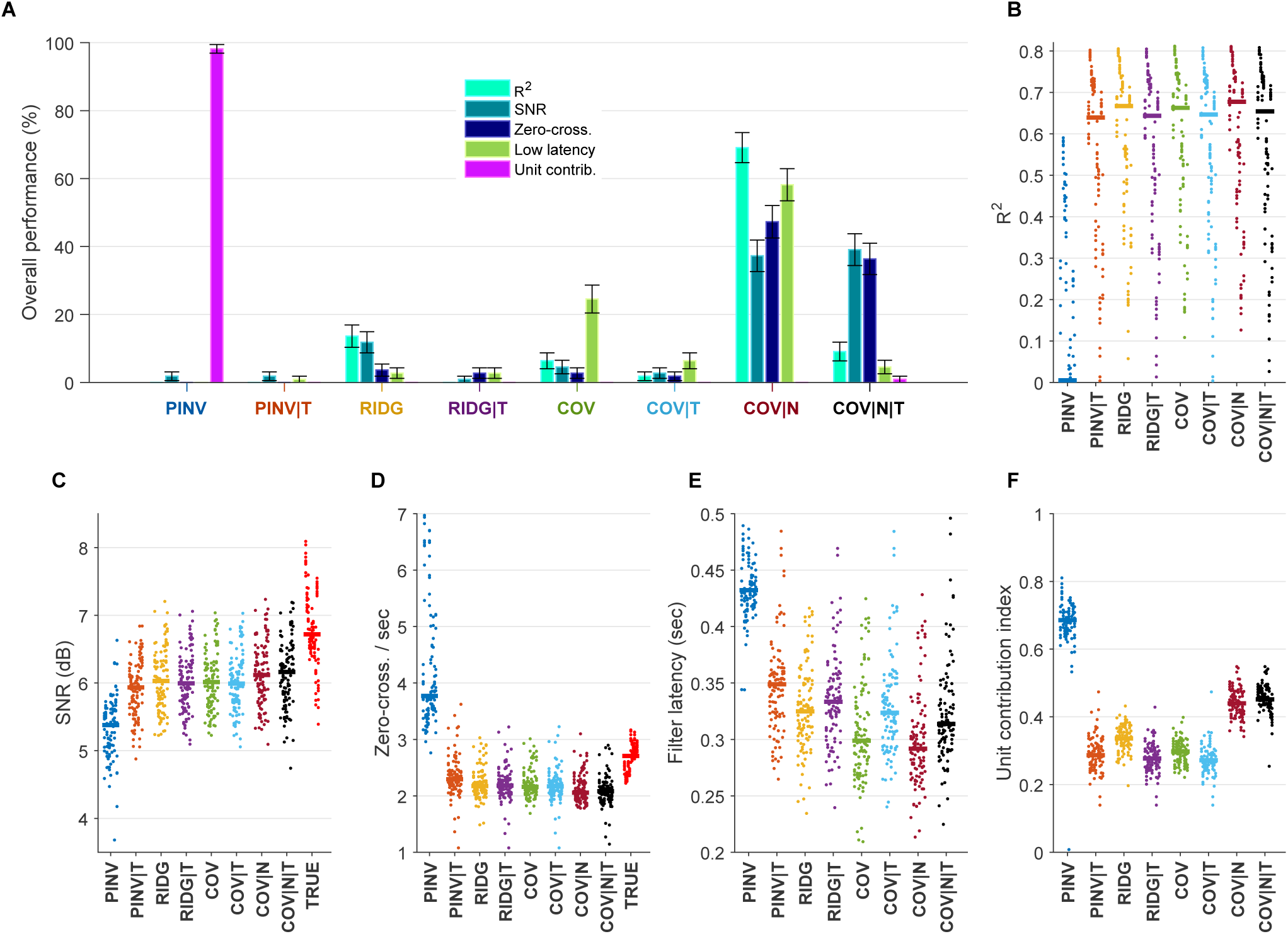
Generalization performance of biomimetic decoding algorithms. (**A**) Percentage of block arrangements across all sessions in which a particular algorithm performed better than all other algorithms. Each percentage is computed as the number of times a particular algorithm outperformed all others, normalized by the total number of block arrangements across all sessions. Error bars represent the standard error of the mean. (**B**-**F**) Performance plots as quantified by different performance metrics. Performance metrics on all block arrangements for all sessions are shown. Each ‘dot’ represents performance on one block arrangement, where each session (containing 10 dots) is displayed in one vertical column, and 11 columns representing 11 sessions are grouped together in the same color for each algorithm, with a solid line representing the median of one group. The performance metrics evaluated on the actual hand velocity (TRUE) in the same block arrangements is also displayed where applicable (red color). In all panels, statistical significance for the PINV algorithm was assessed using Wilcoxon signed rank test with Bonferroni correction for multiple comparisons. (**B**) Decoding performance as measured by the coefficient of determination, *R*^2^, where the PINV algorithm performed significantly worse than all other algorithms (*p <* 0.05), and in some cases had negative *R*^2^ values (trimmed in the current plot). (**C**) Decoding performance as measured by SNR, where the PINV algorithm performed significantly worse than all other algorithms (*p <* 0.05). (**D**) Average decode zero-crossings per second, where the PINV algorithm had significantly higher values than all other algorithms (*p <* 0.05). (**E**) Decoding performance as measured by decoder filters latency, where the PINV algorithm performed significantly worse than all other algorithms (*p <* 0.05). (**F**) Unit contribution indices for decode computation in all algorithms, where PINV had the highest values than all other algorithms (*p <* 0.05).

We also used the same statistical test to study the effect of ‘truncation’ on all algorithms. Truncation simultaneously improved PINV and worsened all others in terms of *R*^2^ (*p* < 0.05), possibly due to the suboptimal greedy algorithm for tuning *M* and *μ*^2^. The same effect was observed in terms of SNR (*p* < 0.05), except for COV|N|T (not significant). For the zero-crossings per second, truncation significantly reduced this metric for PINV only (*p* < 0.05), whereas the result is not significant in all others. Truncation significantly increased decoder filters latency in all algorithms (*p* < 0.05), except for RIDG (not significant). Finally, truncation significantly decreased unit contribution index of all algorithms (*p* < 0.05), except for COV|N where the effect was reversed (*p* < 0.05).

Linear decoding of velocity signals involves a strong trade-off between reconstruction of characteristic velocity spikes versus low-amplitude ‘noise’ bands. A linear decoder can be made to better reconstruct velocity spikes simply by applying a ‘decoder gain’ that scales up the decode (**Sussillo et al.**, 2012). However, such a decoder is expected to perform very badly in terms of ‘target hold time’ because the uniformly applied gain also amplifies the ‘noise’ band – making it more difficult to ‘stop’ near the target (**Sussillo et al.**, 2012; **Gowda et al.**, 2012, 2014; **Golub et al.**, 2014; **Marathe and Taylor**, 2015). This phenomenon can be seen in the representative examples in Supplementary Figure 4, where PINV solution was able to reconstruct a negative velocity spike at the expense of amplifying the ‘noise’ bands surrounding this velocity spike. Regularized solutions generally do *not* suffer from this effect, since the regularization parameter can be tuned to find the best compromise. Figure 5 examines this trade-off from a different perspective. For each algorithm, the block arrangement that produced highest decode SNR on test data was selected, and the average velocity spikes are shown for that block arrangement. These average velocity spikes only represent the ‘signal’ – as opposed to ‘noise’ – component. Even though PINV almost reproduced the average velocity spikes of the actual hand kinematics, its decode has a relatively lower SNR – indicating that this algorithm is also amplifying the ‘noise’ component. Additionally, whereas both RIDG and COV|N produce comparable decode SNR, they achieve this result by two different mechanisms – as can be seen from the respective average velocity peaks – where RIDG puts more emphasis on ‘noise’ suppression and COV|N emphasizes the ‘signal’ component more.

**Figure 5.**
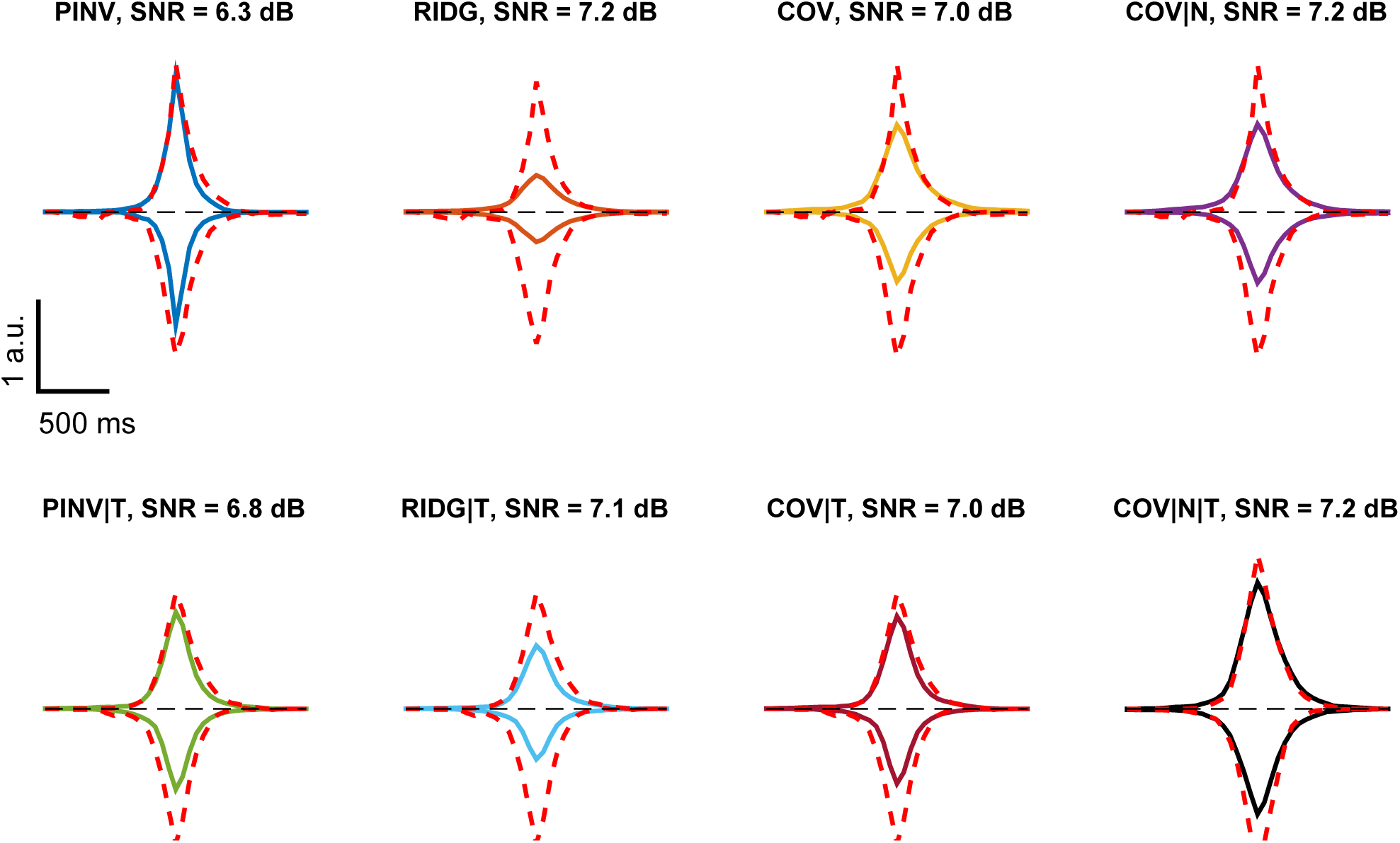
Representative average velocity spikes. Average velocity spikes corresponding to test blocks with best SNR for respective biomimetic decoding algorithms (solid lines, different colors for different algorithms), plotted against average velocity spikes of the natural hand velocity signal taken from the same blocks (dashed red lines). Dashed black lines represent zero baseline. Average positive (correspondingly, negative) velocity peaks are shown above (correspondingly, below) the zero baseline.

Supplementary Figure 5 demonstrates the spatio-temporal structure of decoder filters, and echoes the same observations related to sampling from prior distributions as was shown in Figure 2.

#### 3.3 SINGLE-UNIT DYNAMICS AND UNIT FILTER FUNCTIONS

To provide more evidence for our ‘matched filter’ hypothesis, we calculated single-unit PETHs as described in subsection 2.8. The event time we used in constructing the PETHs was the time of the positive velocity peak (global maximum) of each trial. We then calculated the unit-filter match magnitude and unit fraction metrics. Whereas decoder filters are constructed from limited training data, PETHs are constructed from all trials in a given session. In most units, we found significant correlations (*p <* 0.05) between a unit filter function and the corresponding unit PETH. One representative example is shown in Figure 6. By pooling performance data across all algorithms, we investigated the correlation between different performance metrics introduced in subsection 2.9 and the strength of unit-filter matching (Figure 7). We found highly significant correlations (*p* < 10^*-*10^) mostly in the direction of better decoding performance with higher unit-filter matching strength.

**Figure 6.**
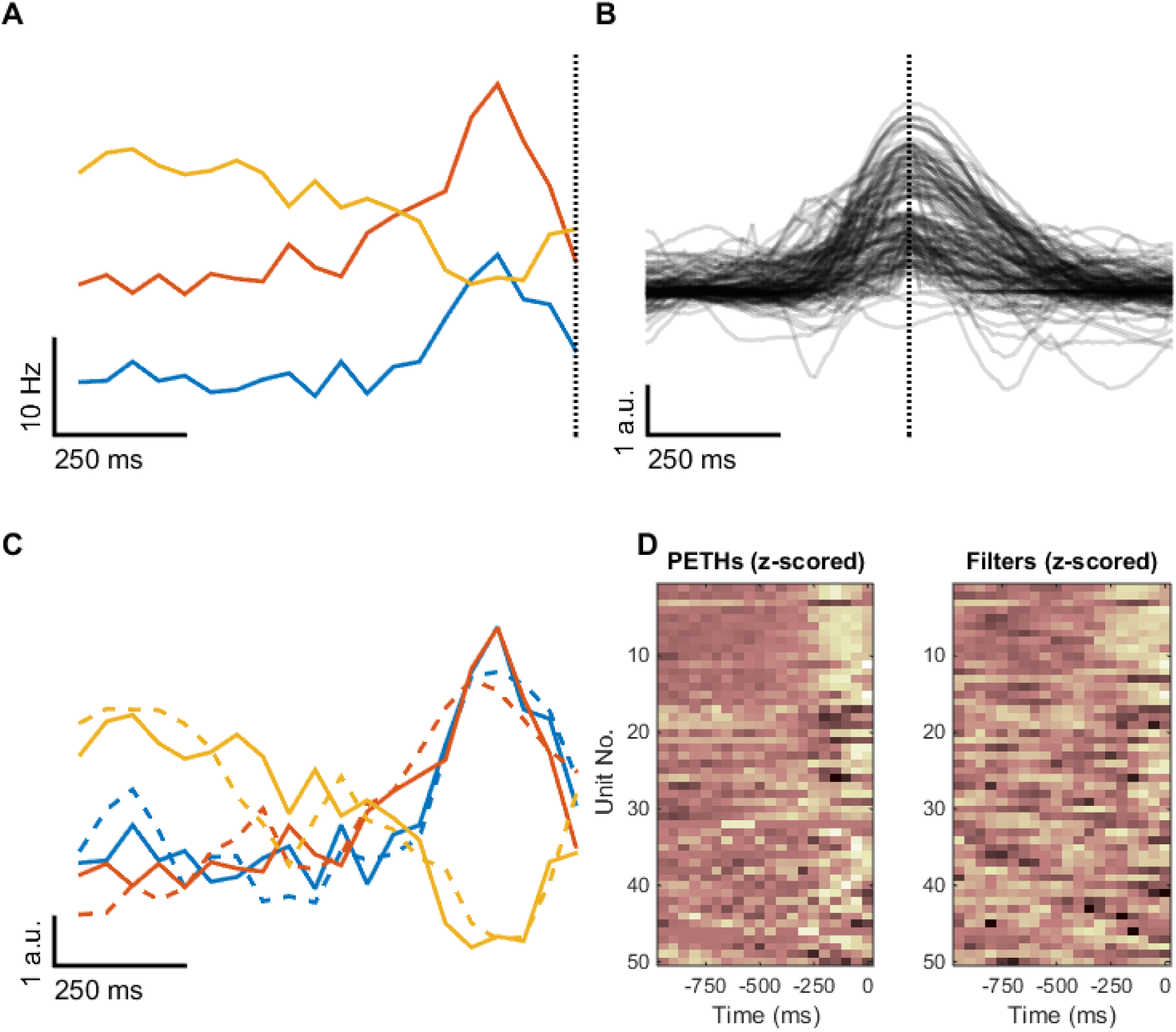
Correlation between single-unit PETHs and corresponding unit filters. (**A**) Top-3 PETHs that correlate with corresponding unit filter shapes in a representative session with a unit-filter match magnitude greater than 0.5. PETHs are constructed relative to the time of the peak events in **B** (dashed line). (**B**) Representative positive velocity spikes extracted from the trials of the same session as in **A**. Dashed line marks the global velocity peak of each velocity spike. (**C**) Visualization of the top-3 PETHs (z-scored) from **B** overlaid with the corresponding z-scored, time-reversed, unit filter functions (dashed lines). (**D**) Visualization of the top-50 correlated PETHs and unit filters (both are z-scored).

**Figure 7.**
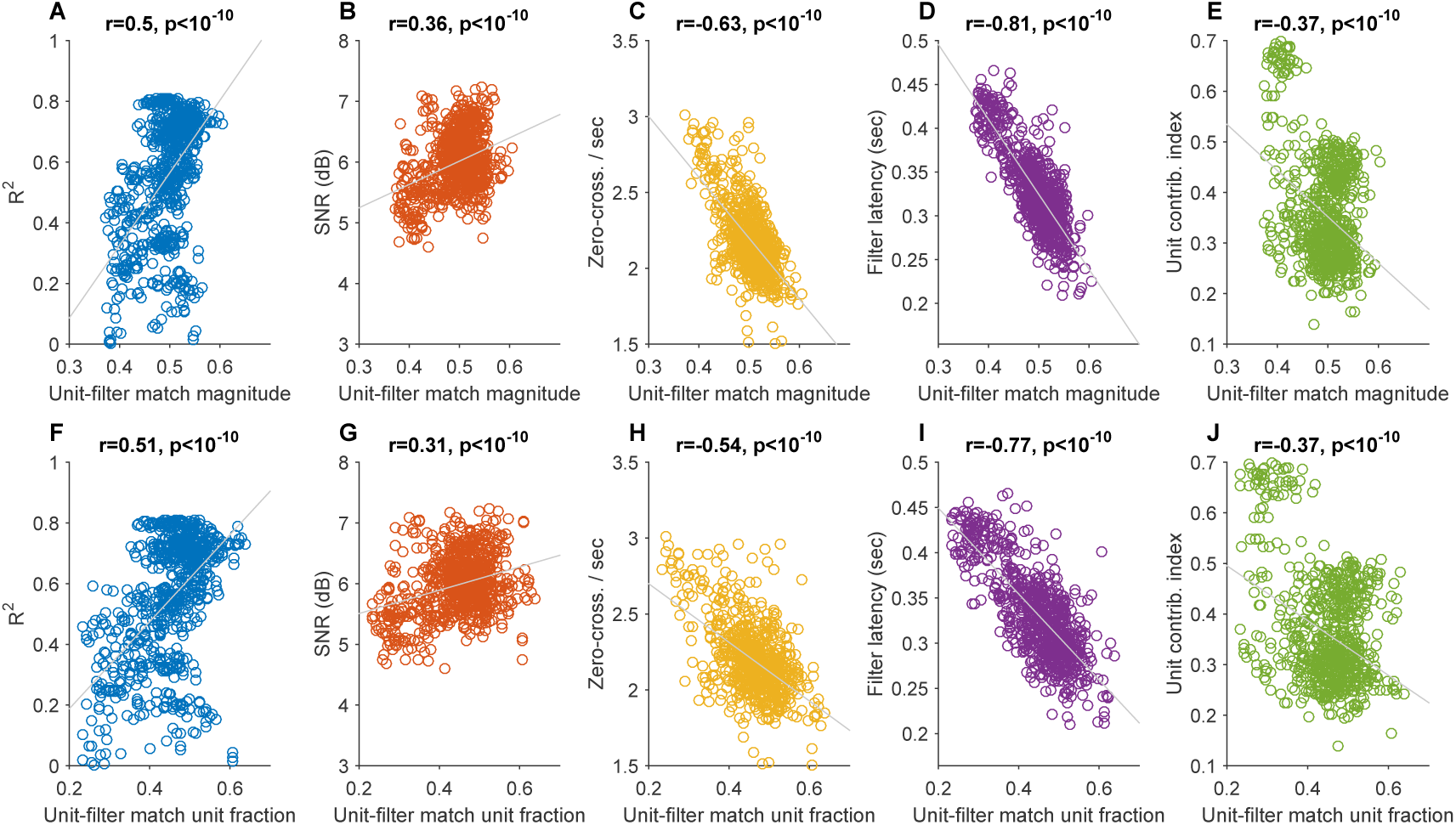
Relationship between unit-filter match metrics and other decoding performance metrics. (**A**–**E**) Relationship between unit-filter match magnitude and decoding performance metrics for all block arrangements from all algorithms after excluding outliers. Outliers in each metric were excluded using the same criterion used in box plots to detect outliers, which relies on interquantile ranges. (**F**–**J**) Same for unit-filter match unit fraction.

#### 3.4 EFFECT OF DECODING PAST AND FUTURE MOVEMENTS

Training of linear Wiener-style decoders is typically done on synchronized sets of neural and kinematic data where the history of the neural input prior to a specific time *t*, is used to decode *current* kinematic signal at the same time *t*. This problem is known in estimation theory as the *filtering* problem. However, it is also possible to decode *future* kinematic signals at times *t* + *k*, with positive *k* – which is known as the *prediction* problem – and to decode *past* kinematic signals at times *t – k* – which is known as the *smoothing* problem (**Wiener**, 1949). We analyzed the decoding performance as it relates to decoding of past, current, and future data. We trained regularized decoders (RIDG, COV, and COV|N) to decode past, current, and future data by artificially changing the temporal alignment of the velocity signal. Decoding future (past) data amounts to moving the velocity signal *k* bins backward (forward) in time. Figure 8 shows the results. We also examined the decoder filters structure and their relation to unit PETHs constructed using the same time shifts in the velocity signal. Figure 9 demonstrates one representative example. This effect is also quantified in Figure 8 using the filter half-RMS time *t*_0.5_.

**Figure 8.**
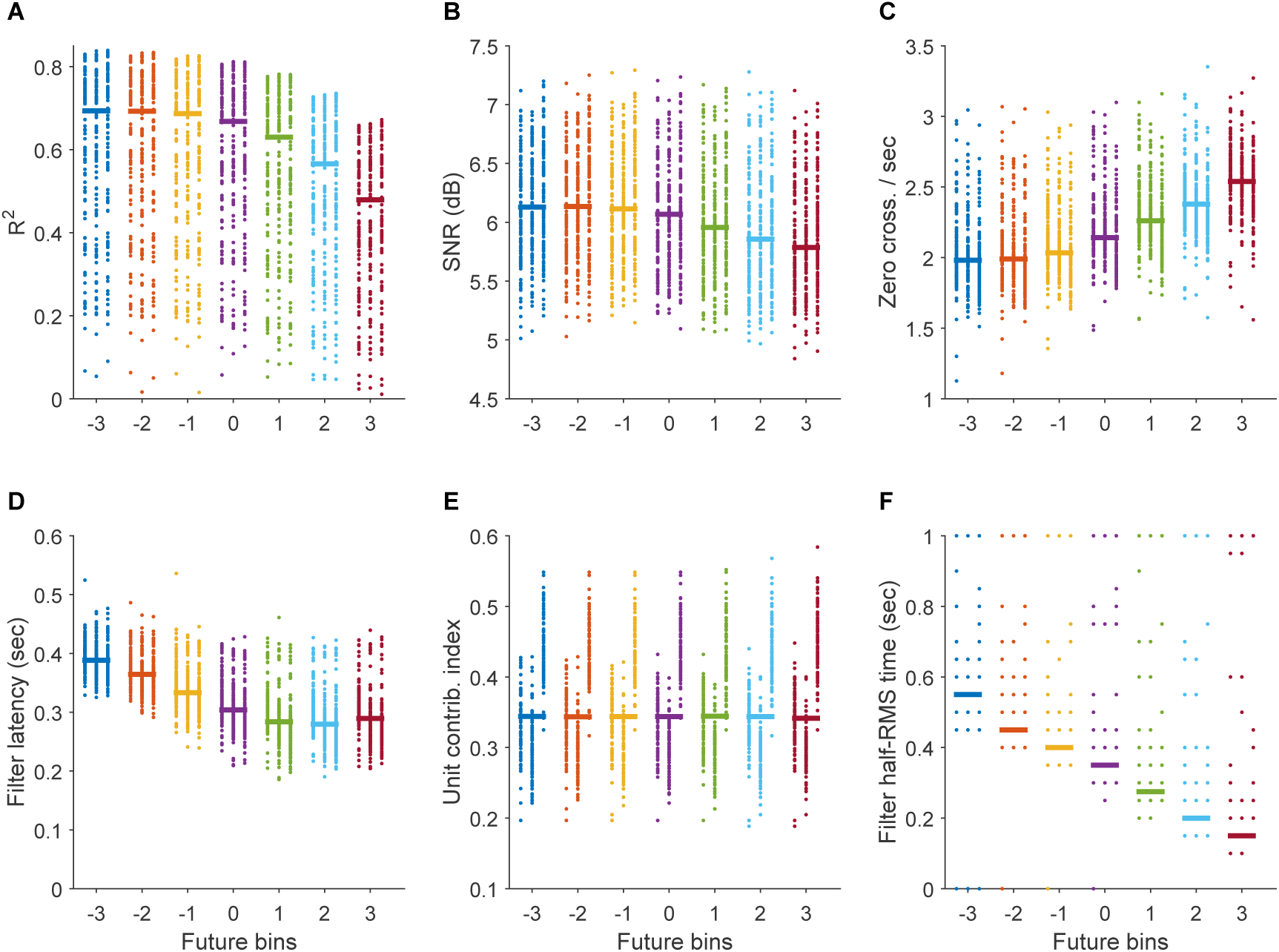
Effects of decoding past and future data on decoding performance and filters structure. Effect of decoding past and future data on: *R*^2^ (**A**), SNR (**B**), zero-crossings per second (**C**), decoder filter latency (**D**), unit contribution index (**E**), and filter half-RMS time (**F**). In all cases, at each future bin value (in the range [-3,3]), each column shows results on test data from *all* 110 block arrangements corresponding to (from left to right) RIDG, COV, and COV|N kernels.

To determine the statistical significance of the trends seen in Figure 8, we pooled the data from the three algorithms and used Wilcoxon signed rank test with Bonferroni correction for multiple comparisons. We found that with increasing *k* the *R*^2^ metric significantly decreased (*p* < 0.01), the zero-crossings per second significantly increased (*p* < 0.01), and the filter half-RMS time *t*_0.5_ significantly decreased. The SNR significantly decreased only for *k ∈* [*-*1, 3] (*p <* 0.01), and did *not* significantly change for *k ∈* [*-*3, -1] (*p <* 0.01). This means that the SNR deteriorates with decoding future data, improves with decoding past data for one bin only, and does not improve with decoding further bins in the past. Filter latency of the resultant decoders significantly decreased with increasing *k* up to +2 (*p <* 0.01). However, it significantly increased with increasing *k* from +2 to +3 (*p <* 0.01). The unit contribution index did *not* significantly change with increasing *k* up to +2 (*p <* 0.01, double-sided), and it significantly increased with increasing *k* from +2 to +3.

#### 3.5 NON-BIOMIMETIC DECODING PERFORMANCE

Lastly, we used the same data set to study generalization performance of non-biomimetic decoders with one DOF. We partitioned the neural data from each session in block arrangements similar to Supplementary Figure 2 – albeit with five blocks for training data and the remaining five for test data ^9^. We compared the following kernels: the truncated neural covariance kernel (COV|T) and its windowed version (COV|T|W), and the truncated diagonal-normalized kernel (COV|N|T) and its windowed version (COV|N|T|W). For windowing the basis functions of the kernels, we used *t*_0.5_ = 0.35 sec as determined empirically from biomimetic decoders in Figure 8. For truncation of the kernels, i.e. selection of which basis functions to retain, we used all basis functions with output eigenmodes that have zero-crossings per second less than a predefined threshold *ζ*_0_ = 3.5 sec^*-*1^ based on Figure 4. This selection automatically satisfies the corresponding constraint in the optimization problem (55). To solve this constrained optimization, we employed a two-step approach to avoid local optima. The first step is Monte Carlo sampling from the priors (see subsection 3.1. The decode produced from these Monte Carlo decoders are then checked against the Γ *≥* Γ_0_ constraint (with Γ_0_ = 2.5 as determined empirically from natural reach trials), and the decode with the highest SNR is subsequently selected to initialize the constrained optimization solver in MATLAB R2014b (MathWorks, Inc.) to find the final decoder.

As a baseline, we compare these non-biomimetic decoders to heuristic decoders designed using half-second moving-average filters (**Moritz et al.**, 2008; **Moritz and Fetz**, 2011; **Koralek et al.**, 2012; **Clancy et al.**, 2014). Decoded units are partitioned into two partitions contributing positive and negative velocity spikes. The decode is computed by subtracting the average firing rate of the negative partition units from that of the positive partition units. For this scheme to produce nearly symmetric decode, it may be necessary to relatively scale the two averages (**Koralek et al.**, 2012; **Clancy et al.**, 2014). Alternatively, we force the two partitions to have nearly equal firing rate variances. This is known as the partition problem, and there exists a simple greedy algorithm to solve it (**Faigle et al.**, 1989). We sort the units in descending order of firing rate variance, and then assign the unit with the highest variance to the positive partition and the unit with the next highest variance to the negative partition. Each subsequent unit is assigned to the partition that has the least cumulative firing rate variance. Unit filter coefficients are all ones for half a second followed by all zeros for the other half. The filter functions for units in the negative partition are multiplied by -1.

We trained decoders with *C* = 150 units, where this number is selected similar to (**Flint et al.**, 2012). For simplicity, we select these units in each session as the top-*C* units that correlate with actual hand velocity, although we note that a fully non-biomimetic unit selection scheme can be achieved based on the functional connectivity between the units (**Eldawlatly et al.**, 2009). Figure 10 shows the overall decoding performance of non-biomimetic decoders in a format that is similar to Figure 4. Unlike the biomimetic decoding case, there is not a single algorithm that represents the best trade-off between different aspects of decoding performance. We used Wilcoxon signed rank test with Bonferroni correction for multiple comparisons to post-hoc compare algorithmic performance. MA had significantly the worst SNR among all others (*p <* 0.05) with a median of 5.0 dB, and had significantly the highest unit contribution index (*p <* 0.05) with a median of 0.37. MA decode distributions were *not* significantly more symmetric than all others (*p <* 0.05, double-sided) with a median of 2.54. COV|N|T had significantly the best SNR as compared to all others (*p <* 0.05) with a median of 6.9 dB, and significantly the highest unit contribution index as compared to all other kernels (*p <* 0.05) with a median of 0.33. It also had significantly lower zero-crossings per second only when compared to COV|T|W (*p <* 0.05) with a median of 1.85 sec^*-*1^. Finally, COV|N|T|W had significantly the smallest decoder filters latency compared to all others (*p <* 0.05) with a median of 47 ms, except for COV|T|W (not significant).

We also investigated the effects of windowing on the proposed performance metrics. Windowing did not have a significant effect on the symmetry (*p >* 0.05, double-sided), significantly decreased the SNR (*p <* 0.05), significantly increased the zero-crossings per second (*p <* 0.05), significantly decreased the unit contribution index (*p <* 0.05), and significantly decreased decoders filter latency (*p <* 0.05).

For visual comparison, Supplementary Figure 5 shows representative decode traces from test data for all algorithms and for the natural hand velocity (TRUE). Figure 11 shows representative average positive and negative velocity spikes for all algorithms as compared to natural hand velocity spikes. Notably, the MA decode has the worst SNR and also the ‘widest’ average positive velocity spikes as compared to corresponding actual hand velocity, which may indicate an ‘over-smoothing’ effect due to averaging.

## 4 DISCUSSION

### 4.1 EXTRACTION OF MOTOR CONTROL SIGNALS FROM NEURAL POPULATION DYNAMICS

Optimal linear decoders have been widely used for over a decade in the BMI community (**Serruya et al.**, 2002; **Carmena et al.**, 2003; **Paninski et al.**, 2004; **Patil et al.**, 2004; **Hochberg et al.**, 2006; **Fagg et al.**, 2009; **Suminski et al.**, 2010). A neural decoder can be viewed as an extraction algorithm that derives motor control signals from neural data – where in optimal linear decoding this extraction algorithm is a MISO linear system that takes in neural data as an input and produces the decode as an output. The decode is subsequently used online in closed-loop BMI control typically as a position or velocity motor control signal. We demonstrated that the design procedure of optimal linear decoders is indeed equivalent to system identification (**Ljung**, 1999, 2002). State-of-the-art system identification methods rely on direct estimation of a filter function (**Pillonetto et al.**, 2014). However, this function estimation procedure – when performed on short data records – can lead to an *overfitting* problem, where the estimates overly fit the noise component present in the actual output leading to poor generalization performance (**Pillonetto et al.**, 2014). One way to circumvent this problem is to use regularization methods such as ridge regression, which is recently witnessing more widespread use in the BMI community (**Suminski et al.**, 2010; **Willett et al.**, 2013; **Collinger et al.**, 2013; **Balasubramanian et al.**, 2013; **Willett et al.**, 2014; **Wodlinger et al.**, 2015). We demonstrated that ridge regression is a special case of the general class of function estimation methods known as *kernel methods* (**Pillonetto et al.**, 2014), which also admit a Bayesian interpretation using the Gaussian processes framework (**Rasmussen and Williams**, 2005; **van der Vaart et al.**, 2008; **Pillonetto et al.**, 2014). This Bayesian approach enables the incorporation of prior knowledge in a rigorous probabilistic framework. For optimal linear decoding, one way to utilize prior knowledge is to specify the spatiotemporal structure of the unit filters. Recent studies have shown that motor cortical activity in non-human primates (**Churchland and Shenoy**, 2007; **Churchland et al.**, 2010, 2012; **Shenoy et al.**, 2013) and in humans (**Pandarinath et al.**, 2015) exhibits a temporal oscillatory component that is well-described by a dynamical system, suggesting that these decaying oscillatory dynamics can form a dynamical basis from which more complex waveforms can be generated. In a series of studies, we hypothesized that the unit filters structure – for non-biomimetic decoding – should match the neural population dynamics as captured by the neural population covariances (**Badreldin et al.**, 2013; **Badreldin and Oweiss**, 2014). Here we presented for the first time, to our knowledge, a formal approach that can exploit these neural population dynamics in the context of biomimetic decoding. We demonstrated the superior decoding performance of decoder filters that are based on the neural covariance kernels in comparison to pseudo-inverse, truncated SVD, and ridge regression methods. Together, these results suggest that optimal linear decoders should be designed as ‘matched filters’ that correlate with the neural population dynamical basis set in order to produce high-SNR decode waveforms.

### 4.2 BIOMIMETIC AND NON-BIOMIMETIC DECODING PERFORMANCE

Offline decoding performance have long been quantified in terms of how well the decode reconstructs actual kinematic or kinetic data (Carmena et al., 2003; **Patil et al.**, 2004; **Fagg et al.**, 2009; **Koyama et al.**, 2010). However, performance improvements quantified by this metric does not necessarily lead to performance improvements in closed-loop online BMI control (**Koyama et al.**, 2010). Here, we proposed five additional performance metrics posed from the perspective of online control. All these metrics are designed to be independent of the *scale*, or gain, of the decode. In fact, since the gain term can typically be tuned online for BMI control (**Sussillo et al.**, 2012; **Marathe and Taylor**, 2015), we attempt to directly quantify the characteristic shapes and dynamics of both the decoder and the decode – which does not change with a varying gain term.

The first metric we proposed is the SNR of a decode. This metric quantifies how well the velocity ‘signal’ component is separable from the ‘noise’ component. High SNR leads to better noise suppression, and consequently to better ‘stopping’ ability. In fact, to improve the performance of Kalman filter decoders, a recent study suggested dampening the decoded speed in order to improve its ‘stopping’ ability represented by the target hold time (**Golub et al.**, 2014). This strategy is similar to *dead-band suppression* – in which velocity values that are below a predefined threshold are suppressed in order to avoid coupling tiny velocity fluctuations to robotic control (**Balasubramanian et al.**, 2013; **Badreldin et al.**, 2013; **Badreldin and Oweiss**, 2014). Our analysis revealed that the SNR typically drops with linear decoding, where the median drop with the pseudo-inverse algorithm was 1.5 dB, and the median drop range for regularized linear decoders was 0.8–1.0 dB. The smallest SNR drop occurred with the diagonal-normalized neural covariance kernels (see supplementary Figure 7), which suggests that designing unit filters as matched filters may improve the decode SNR.

The second metric we proposed is a decode’s average zero-crossings per second. We showed that the median increase in this metric due to linear decoding with the pseudo-inverse algorithm was 1.09 sec^*-*1^ with a maximum of 4.79 sec^*-*1^, indicating that these decoders produce more noisy outputs. On the other hand, this metric decreased with regularized linear decoders, with a median drop range of 0.34–0.57 sec^*-*1^ (see supplementary Figure 7).

The third metric quantifies the average delay, or filter latency, associated with a given decoder. This filter latency is a major contributor to feedforward delays in a closed-loop control system, and it can adversely affect BMI control (**Willett et al.**, 2013; **Marathe and Taylor**, 2015). We showed that the pseudo-inverse decoders have the highest latency with a median of 430 ms, and the diagonal-normalized neural covariance kernels produce decoders with the lowest latency with a median of 290 ms.

The fourth metric is a unit contribution index which quantifies the number of units that contribute more than 90% of the overall magnitude of all unit outputs. This fourth metric is posed from a pragmatic standpoint to minimize the impact of unit loss on decoding performance (**Heliot et al.**, 2010; **Eleryan et al.**, 2014). We showed that among all regularized algorithms, the diagonal-normalized neural covariance kernel performed best in terms of this metric.

The fifth metric – which we mainly used for the non-biomimetic decoder design – is concerned with the degree of symmetry in the distribution of a decode. Most BMI tasks involve some form of symmetry in the task space, and this metric ensures that non-biomimetic decoders can still span the task space as efficiently as their biomimetic peers.

These five proposed metrics are not only scale-invariant, but they are also agnostic of the actual kinematic signals – making them particularly suited to assess the performance of non-biomimetic decoders and compare them to their biomimetic counterparts. On the one hand, non-biomimetic decoding using the moving average filters had the worst decrease in SNR, with a median of 1.8 dB. On the other hand, kernel-based non-biomimetic decoders did not suffer as much drop in SNR, with a median range of 0.1–0.6 dB, thereby exceeding their biomimetic peers for which the median SNR drop range was 0.8–1.0 dB (see Supplementary Figure 7). Additionally, the windowing approach we used to ‘taper’ the basis functions of the proposed kernels resulted in a great reduction of the decoder filters average latency by about ten folds – without much reduction in other performance metrics (see Figure 10). With the diagonal-normalized neural covariance kernels having the highest SNR and also highest unit contribution index values, we conclude this analysis by pointing out that these kernels provide a good compromise in terms of the decoding metrics that we used.

### 4.3 OPTIMAL LINEAR DECODERS AS MATCHED FILTERS

The main hypothesis we put forward in this study is that the unit filters of optimal linear decoders should match the spatiotemporal dynamics of the decoded units. To this end, we quantified this ‘matching’ effect using the PETHs of the decoded units. We proposed two metrics based on Pearson correlation coefficients between unit filter functions and corresponding unit PETHs. These metrics are the unit-filter match magnitude – which quantifies the correlation magnitude, and the unit-filter match unit fraction – which quantifies the fraction of units significantly correlated with the filters. Using these two metrics, we demonstrated universal and significant correlations – using pooled data from all biomimetic linear decoders – between improvements in the unit-filter matching and improvements in the proposed decoder performance metrics. In particular, the unit-filter match metrics positively correlated with the coefficient of determination and the SNR, whereas they negatively correlated with the average zero-crossings per second and the filters latency. It is worth noting that the PETHs were computed using all data on a given session, whereas linear decoders were only computed using a very limited subset of the data on the same session. Our results suggest that the better the matching between the filters and the PETHs, the better these filters approach optimal performance. Moreover, consistent with a matched filter hypothesis, our results indicate that decoding of past and future data using finite filters is equivalent to the ‘causal’ truncation of these filters ^10^.

The decode SNR improvement with better unit-filter matching provides further evidence that harnessing the neural population dynamics may improve decoding performance. Since the population vector algorithm (**Moran and Schwartz**, 1999) – as well as its optimal linear estimation variant (**Koyama et al.**, 2010) – and the velocity Kalman filter algorithm (**Kim et al.**, 2008, 2011) do not take these neural population dynamics into account, we surmise that performance of kernel-based linear decoders that account for neural dynamics may in general surpass other linear decoding algorithms both in terms of the coefficient of determination as well as the decode SNR.

### 4.4 EFFECT OF DECODING FUTURE MOVEMENT

To our knowledge, there is only one study that have investigated the effect of decoding future movement on BMI performance (**Willett et al.**, 2013). The results presented here provide more insight into the online performance improvement reported therein. It is known that motor cortical activity leads natural movement by approximately 100–200 ms (**Moran and Schwartz**, 1999; **Paninski et al.**, 2004). In fact, visual inspection of the spatiotemporal PETHs presented here and also the spatiotemporal unit tuning plots in (**Paninski et al.**, 2004) confirms that motor cortical activity reaches its maximum around this lead time. Our analysis further demonstrates – in agreement with a ‘matched filter’ hypothesis – that optimal linear decoders coefficients also have their maxima around the same lead time (see e.g. Figure 9). Consequently, the effect of decoding future movement on the decoders structure is simply to ‘push’ the decoder coefficients towards immediate neural firing history. This temporal shift in the unit filters structure may also be the main reason for reducing the filters latency (Figure 8). Moreover, the drop in the decoding performance of future data may be explained by comparing the large magnitudes of the decoder coefficients that ‘move out’ of the decoder’s one-second window to the small magnitudes of the coefficients that ‘move in’ into the same window, which also results in the unit filters structure becoming more ‘flat’. In this process, the decoder loses its matched filter structure thereby reducing its decode SNR and – in the extreme case of decoding *k* = +3 future bins – increasing its latency. This observation also resonates well with the deteriorating decode SNR of moving-average decoders (Figure 10 and Supplementary Figure 7) mainly because of their ‘flat’ structure that uniformly weights neural activity across time.

**Figure 9.**
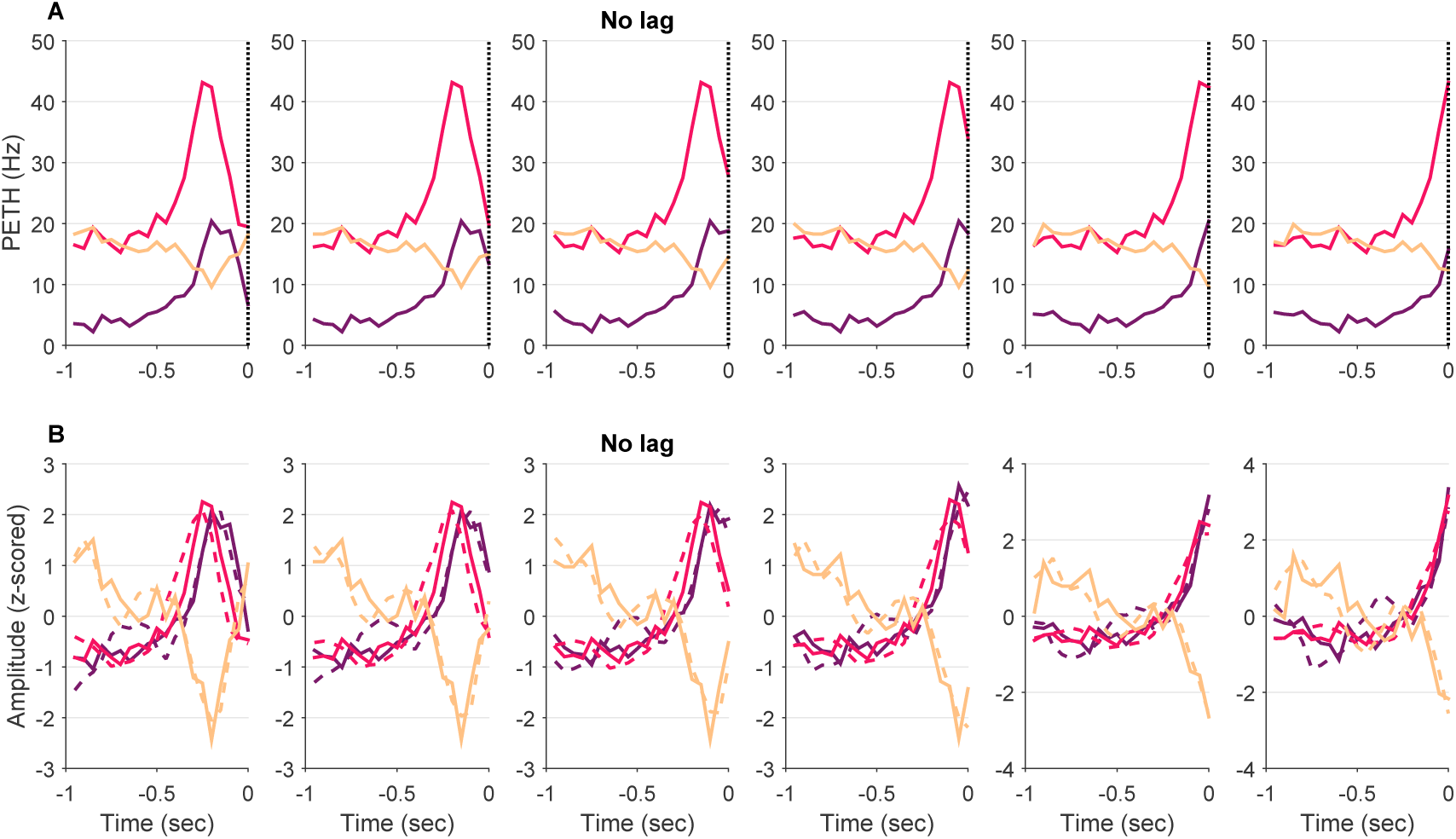
Representative example of changes in the unit-filter match with decoding past and future data. (**A**) Representative PETHs from three units selected as the units with top-3 correlation coefficients between their PETHs and unit filter functions at no lag. The PETHs are constructed for decoding future bins running from -2 (leftmost) to +3 (rightmost). Different colors represent different units. (**B**) Same PETHs as in **A** after z-scoring (solid lines) for visual comparison of their match to corresponding z-scored unit filter functions (dashed lines). Same color scheme and future bins scheme as in **A**.

**Figure 10.**
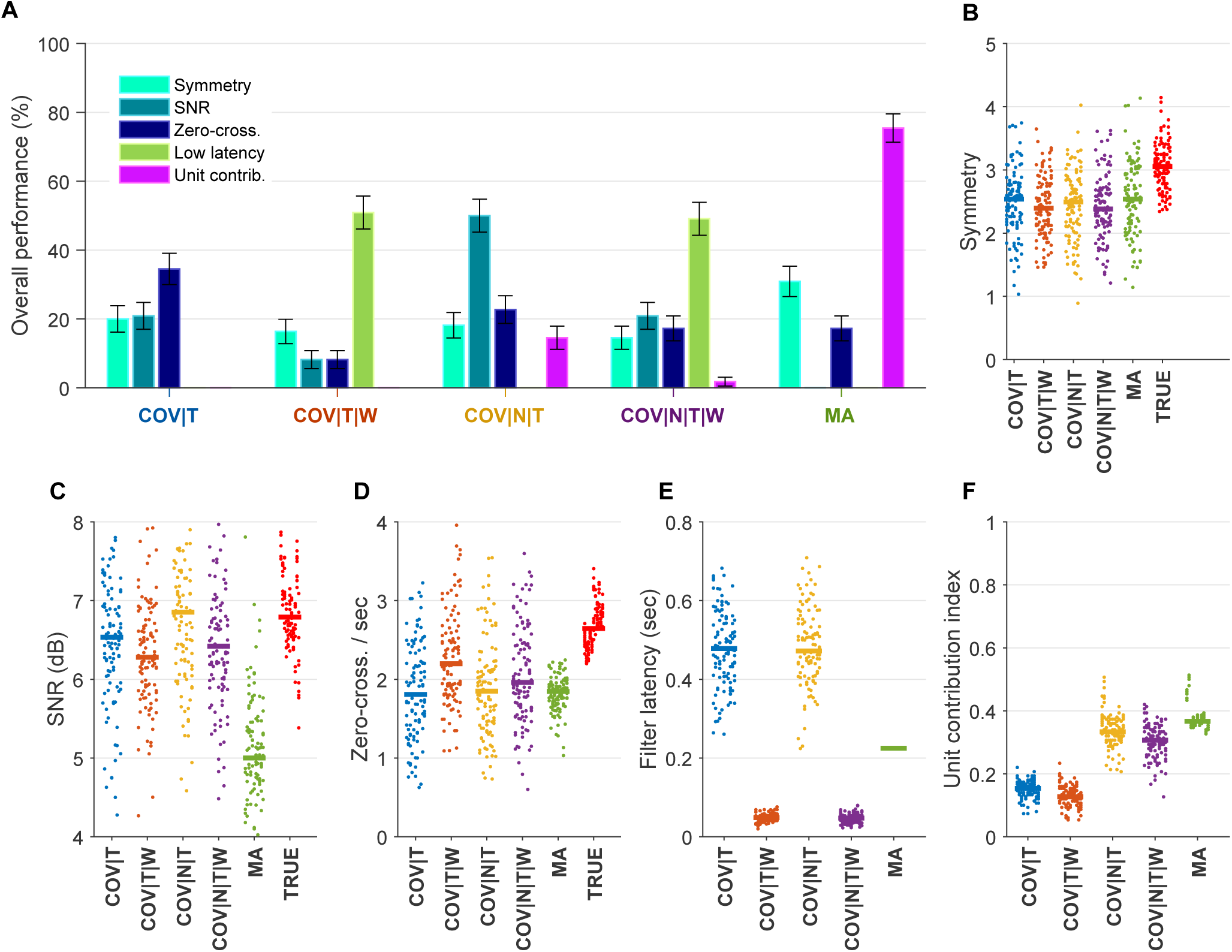
Generalization performance of non-biomimetic decoding algorithms. (**A**) Percentage of block arrangements across all sessions in which a particular algorithm performed better than all other algorithms. Each percentage is computed as the number of times a particular algorithm outperformed all others, normalized by the total number of block arrangements across all sessions. Error bars represent the standard error of the mean. (**B**-**F**) Performance plots as quantified by different performance metrics. Performance metrics on all block arrangements for all sessions are shown. Each ‘dot’ represents performance on one block arrangement, where each session (containing 10 dots) is displayed in one vertical column, and 11 columns representing 11 sessions are grouped together in the same color for each algorithm, with a solid line representing the median of one group. The performance metrics evaluated on the actual hand velocity (TRUE) in the same block arrangements is also displayed where applicable (red color). (**B**) Decoding performance as measured by the decode skewness (or asymmetry) metric. (**C**) Decoding performance as measured by SNR, where the MA algorithm performed significantly worse than all other algorithms (*p <* 0.05; Wilcoxon signed rank test with Bonferroni correction for multiple comparisons) (**D**) Average decode zero-crossings per second. (**E**) Decoding performance as measured by decoder filters latency. (**F**) Unit contribution indices for decode computation in all algorithms.

**Figure 11.**
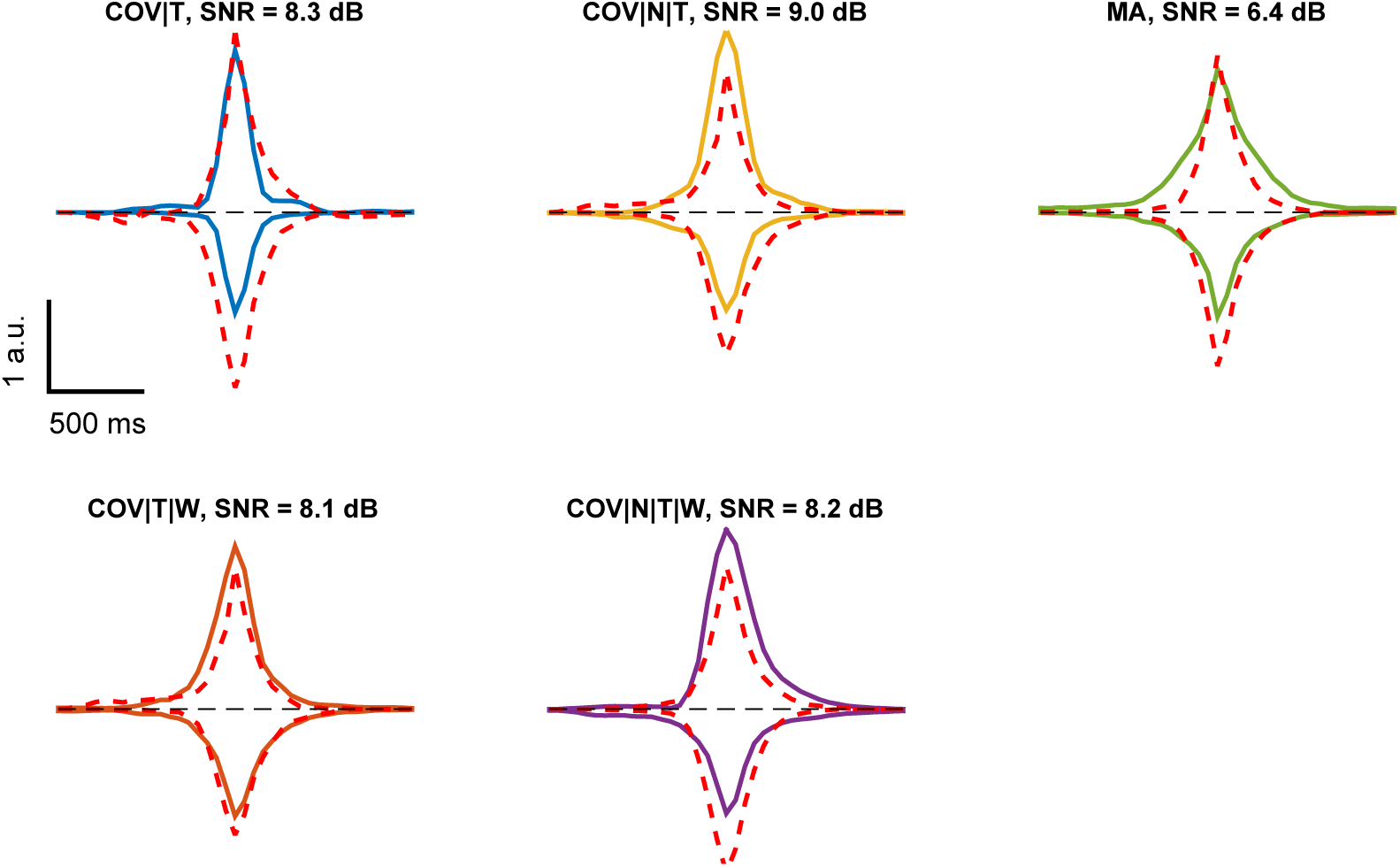
Representative average velocity spikes. Average velocity spikes corresponding to test blocks with best SNR for respective non-biomimetic decoding algorithms (solid lines, different colors for different algorithms), plotted against average velocity spikes of the natural hand velocity signal taken from the same blocks (dashed red lines). Dashed black lines represent zero baseline. Average positive (correspondingly, negative) velocity peaks are shown above (correspondingly, below) the zero baseline.

These insights also justify our non-biomimetic decoder design. The neural covariance kernels ensure the necessary ‘match’ between the unit filter structures and the corresponding unit dynamics. Further-more, ‘tapering’ of the kernel basis filter functions promotes decoders with low filter latency. These factors, together with the symmetry of the decode, are expected to lead to improved online BMI control performance.

## 5 CONCLUSIONS AND FUTURE DIRECTIONS

This study presented, for the first time, a unified framework for design of both biomimetic and non-biomimetic decoders. Through the proposed scale-invariant decoding performance metrics, we showed advantages of kernel-based system identification methods for decoder design. Moreover, we provided theoretical and data-based justifications for designing unit filters from a ‘matched filter’ standpoint. We also investigated the effect of decoding future data on the unit filters structure and the decoding performance. We showed that decoding future data can result in lower decoder latency mainly due to the temporal shift in the decoder weights in a manner that is consistent with a ‘matched filter’ hypothesis. This temporal shift in the decoder structure with decoding future data resulted in lower decoder latency at the expense of deterioration in the coefficient of determination and the SNR. Another approach to reduce average filter latency without much deterioration in the SNR was demonstrated using non-biomimetic decoding with synthetically ‘tapered’ filter basis functions. The extent to which improvements in the proposed decoding performance metrics (e.g. filter latency and SNR) can result in improvements in online BMI control remains to be tested in a future study. Our decoder design approach relies on the estimation of the neural covariances from data and then using these covariances as a kernel. Therefore, we posit that Bayesian covariance estimation could outperform the ML covariance estimation used here – and result in even more decoding performance improvements with short data records. This hypothesis is supported by the optimal kernel theory for system identification (**Pillonetto et al.**, 2014) and can be tested in a future study.

The results presented here are for decoding only one DOF. In biomimetic decoder design, optimal linear decoders for different DOFs are designed simply by optimizing one DOF decoder at a time^11^. However, for non-biomimetic decoders, this is not generally the case. One multi-DOF decoding approach we have used earlier is to partition the pool of recorded units into a number of partitions equal to the number of DOFs based on the unit functional connectivity (**Balasubramanian et al.**, 2013; **Badreldin et al.**, 2013). An alternative approach would be to generalize the theoretical decoder optimization problem (55) to handle multiple DOFs, for example by using the vectorized version of the symmetry metric proposed here. We also note that the theory presented in this study is very abstract and can be extended for the design of infinite-duration neural filters (see **Appendix 1** in the supplementary material) that admit a recursive implementation that is hardware-friendly (**Badreldin and Oweiss**, 2014). Similarly, continuous-time filters can be implemented using mixed-signal integrated circuits to design low-power real-time neural decoding hardware.

In control systems, it is often desirable to obtain an ‘inverse model’ of a plant to be used in a closed-loop control scheme to elicit a desired response from the controlled plant. This ‘inverse model’ – or ‘internal model’ – control scheme can be used for closed-loop neurocontrol by choosing the neurostimulation patterns as outputs of an inverse model of the plant. To this end, the theory we presented is also applicable in this case, where an inverse model of the plant takes the place of a ‘decoder’ (**Liu et al.**, 2011; **Li et al.**, 2013). Finally, kernels based on a finite number of basis functions can be easily adapted online using the least mean squares (LMS) algorithm (**Ninness and Hjalmarsson**, 2005). This is particularly useful for online adaptation in the presence of non-stationaities, for example in closed-loop decoder adaptation (**Orsborn et al.**, 2012) and in the adaptation of the inverse model for control (**Li et al.**, 2013).

## AUTHOR CONTRIBUTIONS

I.S.B. and K.G.O. designed the study; I.S.B. developed the theory and analyzed decoding performance; I.S.B. and K.G.O. wrote the paper.

1 For convenience, the bias term will be dropped throughout this work, without loss of generality, as it can be taken care of by proper offset removal.

2 Also known as Finite Impulse Response (FIR) filters

3 Also known as Infinite Impulse Response (IIR) filters

4 In fact, the decoder filter coefficients estimation problem is a *deconvolution* problem as presented in **Pillonetto et al.** (2014).

5 MIMO decoding using linear filters is handled by cascading multiple MISO decoders.

6 The bin size can be made arbitrarily small, only limited by the highest possible resolution of the sample rate of the raw electrode waveform (Badreldin and Oweiss, 2014)

7 Since the kernel matrix **Q** is positive semidefinite and in general does not have an inverse, the notation ***θ***^*T*^**Q**^*-*1^***θ*** must be understood as presented in **Remark 1** in **Pillonetto et al.** (2014)

8 We make no distinction here between a correlation matrix and a covariance matrix because the mean of the unit firing rates and/or the mean of the decode can be removed online using proper offset removal.

9 The same duration was used for training/test data to facilitate the assessment of the symmetry of the decode between the two data sets.

10 These results are also linked to the orthogonality principle and the mathematical fact that discrete-time filter coefficients at different time bins represent orthogonal spaces.

11 Similarly in linear system identification, MIMO identification is performed as a collection of MISO systems.

## ACKNOWLEDGMENTS

We would like to thank **Flint et al.** (2012) for releasing part of their data to the public, which we used in this study. I.S.B. thanks A. Tamburrino for extensive discussions on the topics of regularization and inverse problems. I.S.B. also thanks J. Southerland, A. H. Fagg, and N. G. Hatsopoulos for stimulating discussions that sparked off the non-biomimetic decoder optimization approach.

*Funding*: This work was supported by NIH-NINDS grant No. NS062031 and DARPA grant No. N66550 001-12-1-4023.

